# Macroscopic Analyses of RNA-Seq Data to Reveal Chromatin Modifications in Aging and Disease

**DOI:** 10.1101/2025.05.14.654062

**Authors:** Achal Mahajan, Francesca Ratti, Ban Wang, Hana El-Samad, James H. Kaufman, Vishrawas Gopalakrishnan

## Abstract

Regulation of gene expression is fundamental for proper cellular function, and is constrained by the local chromatin environment of each gene, which varies spatially along the chromosome and is shaped by epigenetic modifications. Epigenetic modifications induce changes in the local chromatin structure, which can influence gene expression, by affecting the accessibility of DNA to transcription factors. Such changes are particularly relevant in aging and genetic disorders like Hutchinson-Gilford Progeria Syndrome (HGPS) and Werner Syndrome (WRN), where altered chromatin structure contributes to disease pathology. In this study, we analyze RNA-seq data using macroscopic metrics designed to be explicitly sensitive to chromatin modifications. The first metric, intra-chromosomal gene correlation length, measures spatial correlations in gene expressions along the chromosome. The second metric employs an energy landscape model based on the Arrhenius equation to estimate the energetic barriers associated with chromatin state transitions. We apply these metrics to various aging-related datasets, demonstrating their sensitivity to changes in the chromatin structure and the interpretability of the resulting outputs. The intra-chromosomal gene correlation length is particularly effective in quantifying changes in RNA-seq profiles due to increased chromatin accessibility during aging (and conversely, reduced accessibility due to treatment). This metric not only accurately distinguishes cell states, but also provides insight into the direction of aging. For instance, our observations on the effects of anti-sense oligonucleotide (ASO) treatment align with the existing literature, demonstrating that ASO partially restores chromatin structure in diseased cells. They additionally quantify the more pronounced effects in HGPS compared to WRN. The barrier energy landscape further extends this capability by offering a framework for understanding the progressive degradation of the regulatory mechanisms. Together, these metrics provide robust screening tools that enhance our ability to exploit common measurements such as RNA-seq to derive new phenotypes such as chromatin dynamics on aging and disease, offering an alternative perspective that complements traditional analytical techniques and enriches our understanding of cellular states.

## Introduction

The regulation of multiple genes and transcribe-able elements is a fundamental function of the genome, underpinning cellular processes and organismal development (***Busby and Ebrighti, 1994***; ***Walter and Ron, 2011***). Central to this regulation is the local chromatin environment, which exerts a significant influence on gene expression (***Li et al., 2007***). Chromatin structure and its dynamics are critical in regulating these processes, with disruptions in chromatin architecture—such as alterations in TAD boundaries, changes in DNA methylation, and variations in chromatin compaction, are implicated in developmental disorders and cancers (***Sproul et al., 2005***; ***Gerlitz and Bustin, 2011***; ***Hendrich and Bickmore, 2001***; ***Kaiser and Semple, 2017***). Even in senescence, whether natural or pathological, such as in Hutchinson-Gilford progeria syndrome (HGPS) and Werner syn-drome (WRN), chromatin changes like the loss of heterochromatin and telomere shortening have been linked to cellular aging (***Scaffidi and Misteli, 2005***; ***Tsurumi and Li, 2012***; ***Zhang et al., 2015***; ***Lee et al., 2020***; ***Shammas, 2011***). Broadly, the degradation of chromatin structure can lead to significant reorganization of the genome, activating previously silenced genes and contributing to abnormal gene expression and genomic instability (***Csoka et al., 2004***).

The chromatin environment is inherently spatial and varies along the chromosome. Genes within dense heterochromatic regions generally exhibit lower levels of expression than those in more open euchromatin regions (***Huisinga et al., 2006***). In this sense, the chromatin environment constrains transcription. This spatial constraint of transcription by chromatin is modulated by epi-genetic modifications, such as DNA methylation and histone acetylation, which are pivotal in defining the functional and structural distinctions between heterochromatin and euchromatin. These modifications influence gene expression by altering DNA accessibility to transcriptional machinery (***Venkatesh and Workman, 2015***). On a finer scale, nucleosome repositioning differentially regulates genes in response to environmental stimuli or cellular stress (***Jiang and Pugh, 2009***; ***Lai and Pugh, 2017***). For instance, recent studies on human lung and umbilical vein cells have shown that aging cells, with fewer nucleosomes, facilitate faster RNA polymerase II (Pol II) movement, suggesting that structural changes in chromatin can significantly impact transcription precision and error rates (***Debès et al., 2019***). Understanding and quantifying these changes in the chromatin state is, therefore, crucial to advance our understanding of cellular function and dysfunction.

Technological advances over the last two decades have enabled researchers to capture the effects of these changes and understand human biology better, especially with Next-Generation Sequencing (NGS) techniques like RNA-seq, ChIP-seq, and ATAC-seq. Among these different sequencing techniques, the relative ease of performing RNA-seq has made it a standard technology to accompany almost any analysis of the genome approach (***Madsen et al., 2015***), resulting in the creation of large, multi-species databases of prokaryotic and eukaryotic transcriptomes such as Expression Atlas (***Papatheodorou et al., 2018***), Gene Expression Omnibus (***Edgar et al., 2002***), TiGER (***Liu et al., 2008***). Insight mining and computational data analyses on such datasets are typically done using feature extraction and dimension reduction algorithms such as Partial Least Squares (***Datta, 2001***; ***Boulesteix and Strimmer, 2007***), Sliced Inverse Regression (***Bura and Pfeiffer, 2003***; ***Li and Yin, 2008***), principal component analysis (PCA) (***Yeung and Ruzzo, 2001***; ***Lenz et al., 2016***) and singular value decomposition (SVD). These approaches can be useful for identifying diseased phenotypes or genes that are co-regulated and may have similar functions. Although such methods greatly improve our understanding of co-expressing sets of genes involved in a shared biological pathway or process, they do not account for how chromatin properties may influence gene expression (or transcript abundance). This work aims to bridge this gap by describing approaches to illuminate changes in chromatin properties based on transcriptomic data.

Given that chromatin structure creates regions of loosely or tightly packed DNA along a chromosome, affecting transcription, we hypothesize that gene expression will be spatially correlated based on the location of genes within these regions. Specifically, the correlation between neighboring gene expressions is likely to depend on the similarity of their associated chromatin structure and accessibility (***Klemm et al., 2019***). Consequently, genes in the euchromatin domains would exbihit a different correlation pattern compared to the ones in the heterochromatin domains. We note that euchromatin and heterochromatin domains are not necessarily continuous along a chromosome. Instead, the average lengths of these domains determine the rate at which gene expression correlations decay. Loss of heterochromatin in age-related disorders alters chromatin structure, affecting these correlations. This change can serve as an indicator of chromatin state, helping to distinguish diseased states from healthy ones and providing insights into aging and the effects of various perturbations.

While the correlation values act as an indicator variable of the chromatin state, they do not characterize the barrier that the organism/cells need to cross to restore them to a younger state. In other words, heterochromatin-related alterations in DNA caused by histone modifications necessitate energy input in the form of ATP hydrolysis. This process releases energy that the remodeling complexes can utilize to manipulate and modify nucleosomes. Such modifications can potentially impact the accessibility of DNA to transcription factors and RNA polymerase, ultimately leading to changes in gene expression. Getting insight into this energy barrier is crucial for describing the energetics behind the physical processes involved in modifying the nucleosomes on the nanometer scale to individual chromosomes on the micron scale (***Mulligan et al., 2015***; ***Cortini et al., 2016***). Furthermore, the trend on the energy potential across age also would yield insight into the “stress buffer capacity” of the organism at the genomic level. Buffer capacity refers to the ability of biological systems to maintain homeostasis and resist stress or perturbations over time (***Epel and Lithgow, 2014***).

Thus, the objective of the present study is twofold: (1) To develop a data-driven approach to validate the hypothesis that epigenetic modifications, such as those associated with aging, alter chromatin structure and subsequently affect long-range correlations in gene expression; and (2) To characterize the ease of these modifications through thermodynamic principles. Towards these ends, we propose two macroscopic measures that systematically model the effects of aging on chromatin. In proposing a spatially sensitive significant length of correlation between genes on a chromosome (*intra-chromosomal gene correlation length* - *ℓ*^*\**^), we leverage the theory behind the deterioration of the regulatory mechanism due to aging, including loss of heterochromatin and reduced histone density that results in increase in transcription rate and leaky genes (***Debès et al., 2023***; ***Vermulst et al., 2015***; ***Zhang et al., 2021***). This increase in “leaky” genes results in an overall increase in the low-frequency component of gene expressions, resulting in a longer scale of correlations. In modeling the “energy barrier” that cells have to overcome to move from one chromatin state to another, we leverage the principles of transition probabilities, quantifying probabilities of genes jumping ranks between different states (e.g., diseased vs. healthy, old vs. young). These rank jumps reflect the ease with which gene expression levels can shift in response to changes in chromatin structure. We then convert these probabilities into “energy” using the Arrhenius equation, providing a thermodynamic perspective on the stability of these chromatin states. These methods were applied to whole transcriptomic (bulk RNA-seq) datasets from human fibroblasts. Additionally, we demonstrate that these metrics can be utilized both in a standalone setting and dovetailed into a larger solution, leveraging these measures as input features to downstream analytics. Our findings strongly support a link between genome-wide relative gene expression changes and chromatin structure modifications caused by epigenetic alterations.

## Results

### Intra-chromosomal gene correlation length derived from transcriptomic data via spatial autocorrelation

Traditional gene co-expression analyses (***Zhang and Horvath, 2005***; ***Van Dam et al., 2018***) use transcriptomic data to reveal co-expression based on abundance correlations. However, these methods do not probe *spatial* correlations along chromosomes and, consequently, do not reveal changes to chromatin. To address this, we propose a spatially sensitive measure to capture the correlation length of genes along a chromosome. This metric not only provides information about changes in the chromatin structure but also may act as a discriminating feature for other downstream analytics.

Modeling gene-gene correlation to derive the *intra-chromosomal gene correlation length* requires careful consideration, particularly due to non-linear relationships between genes (***Kontio et al., 2020***). We adopt a two-step process, wherein the first step employs Distance Correlation (***Székely et al., 2007***; ***Székely and Rizzo, 2009***) to capture these non-linear correlations. This step quantifies the correlation strength between genes separated by Δ*ℓ* genes, where *ℓ* ranges from 0 to *n* − 1 for a chromosome with *n* genes (see Intra-chromosomal gene correlation length in Methods). The relative positions of the genes define the “neighbor” distance (Δ*ℓ*), where Δ*ℓ* = 0 corresponds to the same gene, Δ*ℓ* = 1 indicates adjacent genes, Δ*ℓ* = 2 refers to next-nearest neighbors, and so on. Gene pairs are grouped into bins based on Δ*ℓ*, and Distance Correlation (𝒞 (Δ*ℓ*)) is calculated for each bin to represent the overall correlation strength between the genes separated by Δ*ℓ*.

In the second step, we calculate the length of this correlation by performing autocorrelation analysis on (𝒞 (Δ*ℓ*)) over different lags (Δ*ℓ*). The autocorrelation, ℛ (*ℓ*), indicates whether genes separated by Δ*ℓ* − 1 neighbors affect the correlation of genes separated by Δ*ℓ* neighbors. The maximum significant autocorrelation lag determines the *intra-chromosomal gene correlation length* (*ℓ*^*\**^) (see Intra-chromosomal gene correlation length in Methods).

Two publicly available human whole transcriptome bulk RNA-seq datasets are used in this study (see Methods). The first dataset, the Long Interspersed Nuclear Element-1 (LINE-1 or L1) dataset (***Della Valle et al., 2022***), comprises human fibroblast samples from 7 cell lines, each with 3 replicates; additional details are provided in ***Table 1***. The second dataset, the Fleischer (FL) dataset (***Fleischer et al., 2018***), includes human fibroblast cells from 143 donors of varying ages, ranging from newborns to 96 years. These datasets were selected based on the observation that both natural aging and Progeria related disorders, such as Hutchinson Gilford Progeria Syndrome (HGPS) and Werner Syndrome (WRN) induce loss of heterochromatin and disrupting transcriptional silencing (***Lee et al., 2020***). Additionally, anti-sense oligonucleotide (ASO) was used as an intervention on HGPS and WRN cell lines in L1 dataset. It was shown that ASO treatment restored hete-rochromain and reversed DNA methylation age (***Della Valle et al., 2022***).

**Table 1.** Cell state definitions in LINE-1. The table outlines the cell states along with their descriptions used in ***Della Valle et al. (2022***), including Wild Type (WT), Hutchinson-Gilford Progeria Syndrome treated with scrambled ASO (HGPS-SCR), untreated HGPS (HGPS-NT), HGPS treated with LINE-1 ASO (HGPS-L1-ASO), Werner Syndrome treated with scrambled ASO (WRN-SCR), untreated Werner Syndrome (WRN-NT), and Werner Syndrome treated with LINE-1 ASO (WRN-L1-ASO)

| Cell State | Description |
| --- | --- |
| WT | Wild Type |
| HGPS-SCR | HGPS treated with scrambled ASO |
| HGPS-NT | HGPS with no treatment |
| HGPS-L1-ASO | HGPS treated with LINE-1 ASO |
| WRN-SCR | WRN treated with scrambled ASO |
| WRN-NT | WRN with no treatment |
| WRN-L1-ASO | WRN treated with LINE-1 ASO |

We observe that gene expression follows a pattern that allows us to categorize genes in the datasets into High Abundance Transcripts (HAT) and Low Abundance Transcripts (LAT) (see Intra-chromosomal gene correlation length in Methods). HAT exhibits a linear trend, while LAT displays a roll-off, possibly due to rarefaction and the presence of genes with very low expression levels (***Supplementary Figure S1***). It is important to note that HAT and LAT are distinguished based on their expression characteristics rather than their association with euchromatin or heterochromatin domains. While it may be intuitive to assume that highly expressed genes originate from euchromatin, this cannot be conclusively stated as a complete representation of euchromatin genes, nor can LAT be definitively linked to heterochromatin.

***Figure 1*** presents the spatial decay of the gene-gene autocorrelation function (ℛ) for the two classes —**(a)** HAT in red and **(b)** LAT in green — using the LINE-1 dataset. The x-axis represents the different lags (*ℓ*). The *intra-chromosomal gene correlation length* (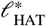 and 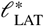) is determined by the point where the autocorrelation falls below the 95% confidence limit (the shaded regions). Observe that 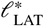 (***Figure 1b***) is longer compared to that of 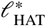 (***Figure 1a***). This is primarily due to the fact that: (1) The LAT gene class is always larger than the HAT class (***Supplementary Figure S1***), and (2) Majority of the genes in the LAT class have similar values (often near 0) (***Figure 2***). These stable and slow-varying expression levels introduce low-frequency components into the system, resulting in longer autocorrelation lags, as gene expression correlations decay more gradually over increasing chromosomal distances.

**Figure 1.**
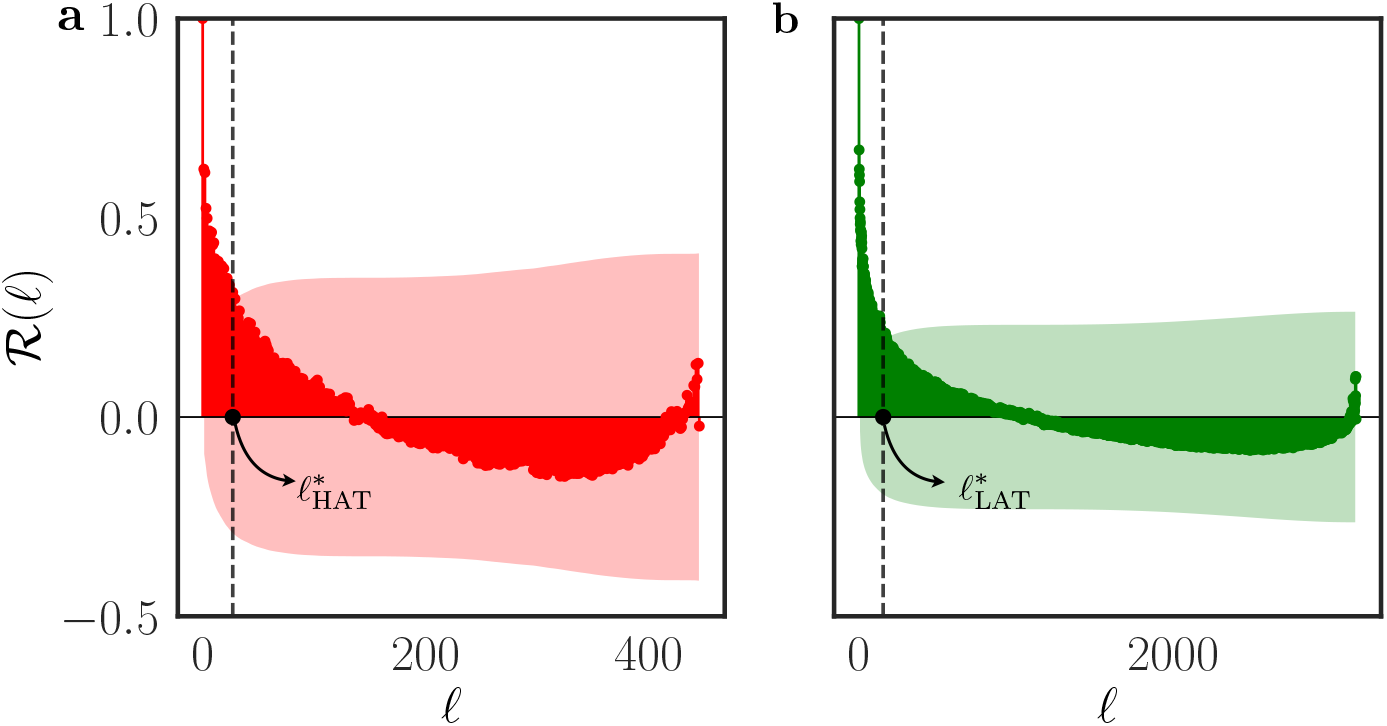
*Intra-chromosomal gene correlation length ℓ*^*\**^ (measured through autocorrelation as a function of lag) for genes in **(a)** HAT (red) and **(b)** LAT (green) class located on chromosome 1. The plots are shown for WT-1 sample in L1 dataset. The *intra-chromosomal gene correlation length* is defined as the point where the autocorrelation falls below the 95% confidence limit. This value is normalized as per the normalization described in Intra-chromosomal gene correlation length in Methods. The correlation length is relatively longer for genes in the LAT class compared to that of genes in the HAT class.

**Figure 2.**
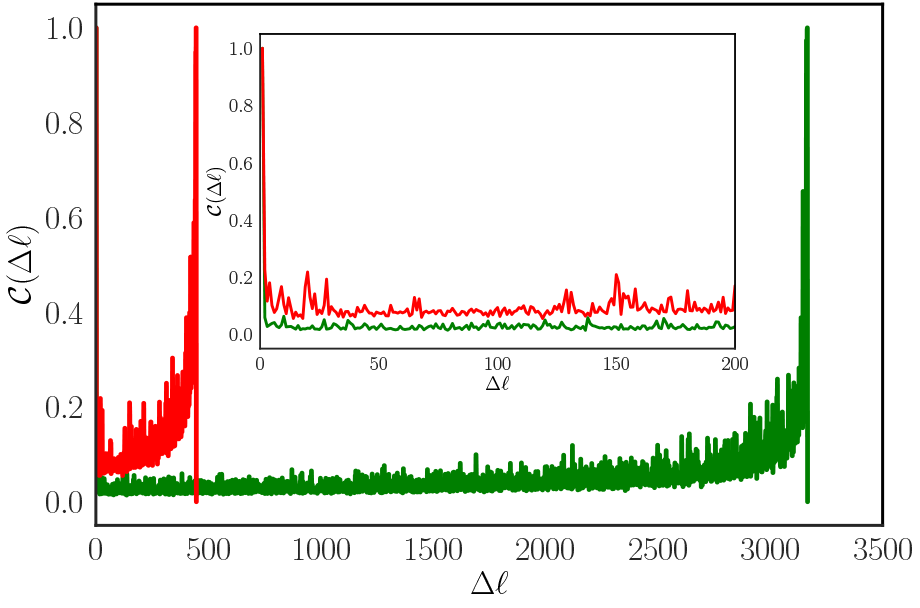
Distance correlation coefficient 𝒞 (Δ*ℓ*) measured based on abundances for binned pairs of genes plotted against Δ*ℓ*, where each bin contains pairs of genes separated by Δ*ℓ*. The red line represents the correlations for HAT class, while green line indicate correlations for the LAT class. The plot depicts data from the WT-1 sample in L1 dataset, focusing on genes located on chromosome 1. The inset figure illustrates the difference in the correlations for a reduced range of x-axis.

***Figure 2*** shows the spatial dependence of the correlation coefficient, 𝒞 (Δ*ℓ*), measured using the distance correlation (***Székely et al., 2007***; ***Székely and Rizzo, 2009***) on the LINE-1 dataset. Given the different ranges of Δ*ℓ* for LAT and HAT, the inset in ***Figure 2*** highlights the range 0 ≤ Δ*ℓ* ≤ 200. For this particular example (chromosome-1 for WT-1), the distance correlation (𝒞 (Δ*ℓ*)) for HAT decays sharply and then fluctuates around a mean value of 0.09. A similar pattern can be seen for genes in the LAT class. It is worthwhile to highlight that the spikes observed at the end of the correlation plot for both high and low transcriptions are artifacts of fewer gene pairs with high Δ*ℓ*.

By definition, the distance correlation is non-negative, with a value of 0 indicating independence between pairs of genes. In our analyses, shown in ***Figure 2***, we observed that the signal from the HAT class is the stronger between two and the signal from the LAT class, being mostly uniform, can be consituted as background noise. Therefore, throughout the remainder of this paper, the *intra-chromosomal gene correlation length* (*ℓ*^*\**^) refers to 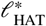 unless stated otherwise.

### Intra-chromosomal gene correlation length serves as a proxy of chromatin structure

Since chromatin structure regulates gene expression, previous studies, such as ***Stavreva and Hager*** (***2015***), have utilized gene expression profiles as proxies to investigate the effects of chromatin states on function. In this work, we build on that concept by examining whether *ℓ*^*\**^ can serve as a lens into chromatin structure. For *ℓ*^*\**^ to function as such, it is essential to verify that this measure: (a) Correlates with gene expression, and (b) Reflects the expected changes in chromatin structure associated with aging.

To assess the level of alignment between (*ℓ*^*\**^) and gene expression trends, we calculated the distance between various cell states and a reference state. For the LINE-1 dataset, we determined the Manhattan similarity distances between different treatment groups and the WT by using normalized gene expression (*α*_*t*_) and (*ℓ*^*\**^) (see Manhattan distance on gene expression and intrachromosomal gene correlation length in Methods). Similarly, in the Fleischer dataset, the 0-20 age group was used as the reference cell state. When two cell states are identical, the distance is 0. A smaller distance indicates greater similarity between the samples. The variation observed within the WT and 0-20 group reflects the inherent variation between replicates, resulting in a relatively lower similarity distance.

***Figure 3*** compares the trend of the distances 𝒟 (*α*_*t*_) and 𝒟 (*ℓ*^*\**^) for LINE-1 and Fleischer datasets. We can see that 𝒟 (*ℓ*^*\**^) exhibits a pattern similar to 𝒟 (*α*_*t*_), indicating that this new proposed measured does capture the functional aspects of gene expression. An interesting observation from ***Figure 3*** is that ASO-treated LINE-1 appears to be more effective in restoring HGPS cell lines closer to wild-type compared to WRN. Another observation from the results on Fleicher dataset is the magnitude of the difference in similarity distance is more pronounced in *ℓ*^*\**^ than in gene expression.

**Figure 3.**
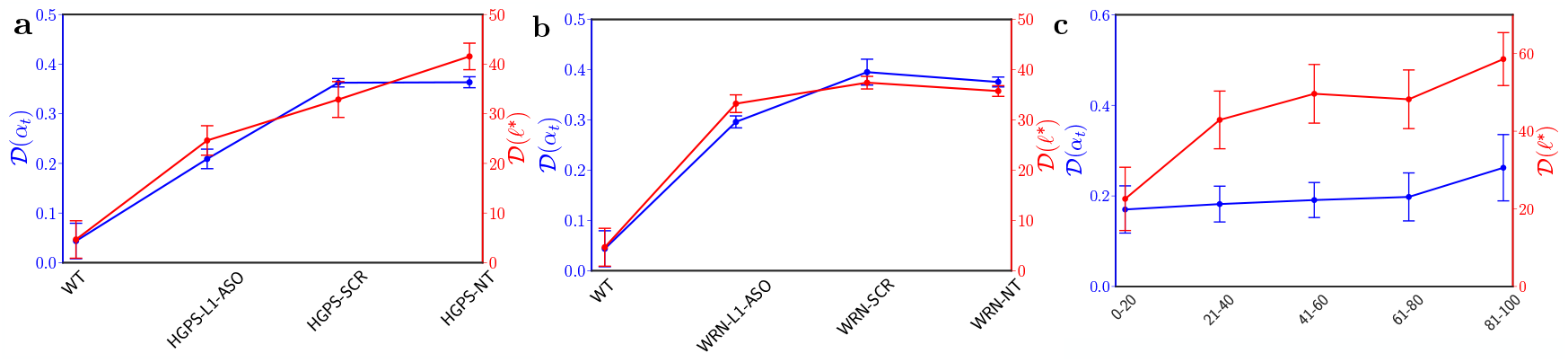
Similarity measured using Manhattan distance for different samples with respect to WT for **(a)** HGPS, **(b)** WRN in L1 dataset and **(c)** Fleicher dataset. The blue and red line represents the similarity on gene expressions and *ℓ*^*\**^ respectively. The plot highlights the distance as measured using *intra-chromosomal gene correlation length ℓ*^*\**^ (red) has a similar trend to the one measured using similarity distance from gene expression *α*_*t*_ (blue).

Now that we have demonstrated that *ℓ*^*\**^ effectively captures similarity distance trends consistent with gene expression patterns, the next question to explore is whether *ℓ*^*\**^, as an indicator of chromatin structure, reveals properties that align with the aging process.

A well-established effect of aging is the loss of heterochromatin. Numerous studies have demonstrated that heterochromatin domains, which are formed early in embryogenesis, deteriorate over time, leading to alterations in global nuclear architecture, derepression of previously silenced genes, and abnormal gene expression patterns (***Villeponteau, 1997***; ***Tsurumi and Li, 2012***; ***Lee et al., 2020***). If total chromosome length remains conserved, the reduction in average heterochromatin domain size would lead to an expansion of euchromatin domains, implying greater accessibility for certain genes. ***Supplementary Figure S2*** illustrates an elevated activation of the gene set in diseased/aged samples from both the LINE-1 and Fleischer datasets. A possible explanation for this phenomenon is the “leaky” gene hypothesis (***Zhang et al., 2021***), where the breakdown of heterochromatin and histone allows transcription of nearby genes by the RNA polymerase. As these “leaky” genes maintain a continuous relatively low level of expression, they result in a steady baseline expression acting as a “low-frequency signal” - signal that changes slowly over time, is smoother and show more sustained pattern. This results in a higher autocorrelation between genes within larger euchromatic domains. As the proposed *intra-chromosomal gene correlation* metric captures this spatial autocorrelation of a gene’s expression, we expect that in aged cells this length (*ℓ*^*\**^) will be longer. Indeed, we observe an increasing trend (Mann Kendall Test, p-value = 0.027, see Methods) in Fleischer dataset, as evident in ***Figure 4a***, although changes in the heterochromatin are not identical for all chromosomes shown by the different degrees of variation of *ℓ*^*\**^ in each age group. When we look at the sample level *ℓ*^*\**^ changes regarding to ages (***Figure 4b***), the increasing trend is still preserved (Mann Kendall Test, p-value = 0.0, see Methods). This observation aligns with the current literature indicating that aging leads to loss of heterochromatin (***Csoka et al., 2004***; ***Scaffidi and Misteli, 2005***; ***Tsurumi and Li, 2012***; ***Zhang et al., 2015***; ***Lee et al., 2020***), and our hypothesis that the proposed metric *ℓ*^*\**^ is sensitive to it. Also, if *ℓ*^*\**^ is indeed capturing the loss of heterochromatin then, not all chromosomes exhibit this erosion at the same scale.

**Figure 4.**
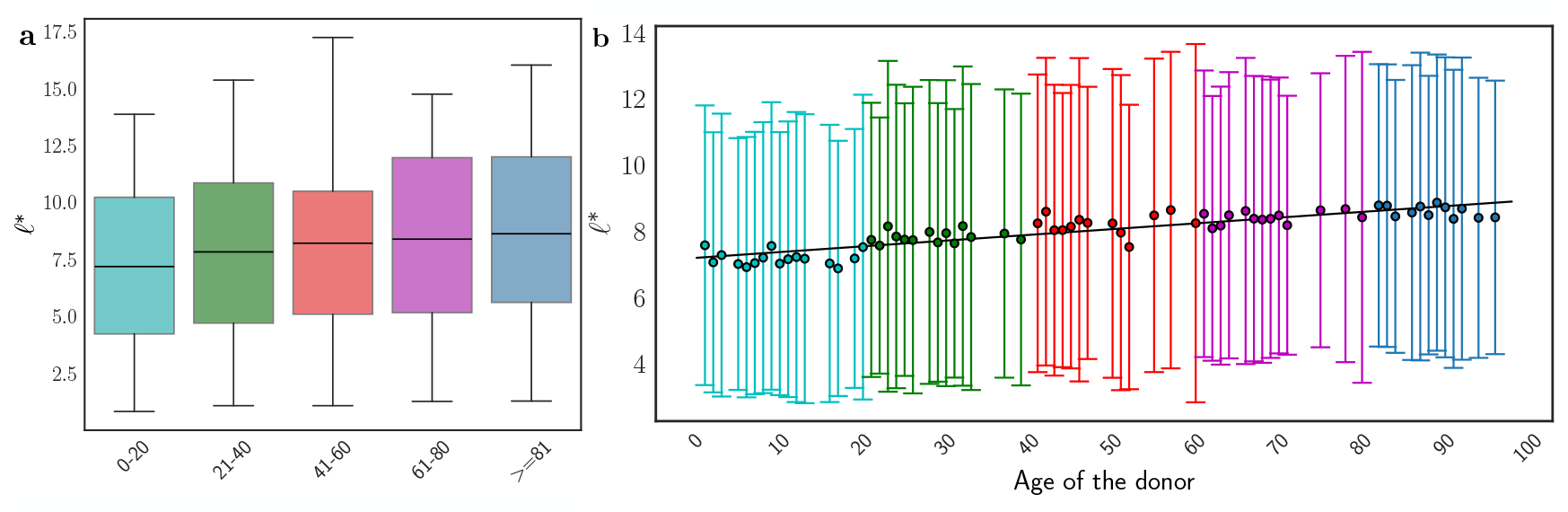
Box plots showing variation of *intra-chromosomal gene correlation length* (*ℓ*^*\**^) across different chromosomes in Fleischer dataset. **(a)** Samples are grouped in age groups ordered from young to aged. Mean of the box plots are labeled. **(b)** *ℓ*^*\**^ for each sample ordered by donor age. The center bar denotes the mean across all chromosomes, and the error bar shows the variation.

However, we did not observe significant ensemble (all chromosomes) level changes in LINE-1 dataset (***Supplementary Figure S5***), possibly indicating a different epigenetic modification mechanism for pathological aging than natural aging. Having established that heterochromatin density varies by chromosome, and consequently *ℓ*^*\**^, we wish to better understand those important chromosomes that are differently expressed across different samples/cell lines and trends of their corresponding *ℓ*^*\**^. Hence, we perform differential gene expression analysis for the (HGPS- and WRN-) NT vs. WT cell lines using PyDESeq2 (***Supplementary Figure S6***, see Methods). In HGPS-NT vs. WT, we observe that the largest number of differentially expressed genes (DEGs) out of the top 200 hits based on the absolute log_2_ fold change are located on chromosome 6 and this chromosome has the highest fraction of DEGs as shown in ***Table 2a***. In WRN-NT vs. WT, chromosome 6 has the second largest number of differentially expressed genes (only 2 fewer than chromosome 1) but still has the highest fraction (***Table 2b***). It is worthwhile to note that the largest cluster of histone genes (*∼* 80%), HIST1, is located on chromosome 6 (***Albig et al., 1997***). Hence, we further investigate the *ℓ*^*\**^ of chromosome 6 as shown in ***Figure 5*** (*ℓ*^*\**^ for all chromosomes is available in the Additional Files category). ASO treatment brings the *ℓ*^*\**^ closer to the wildtype and SCR/NT has the highest *ℓ*^*\**^ values. This effect is more pronounced for HGPS than WRN. For both aging and disease, the data illuminates that each chromosomes should be investigated individually.

**Table 2.**
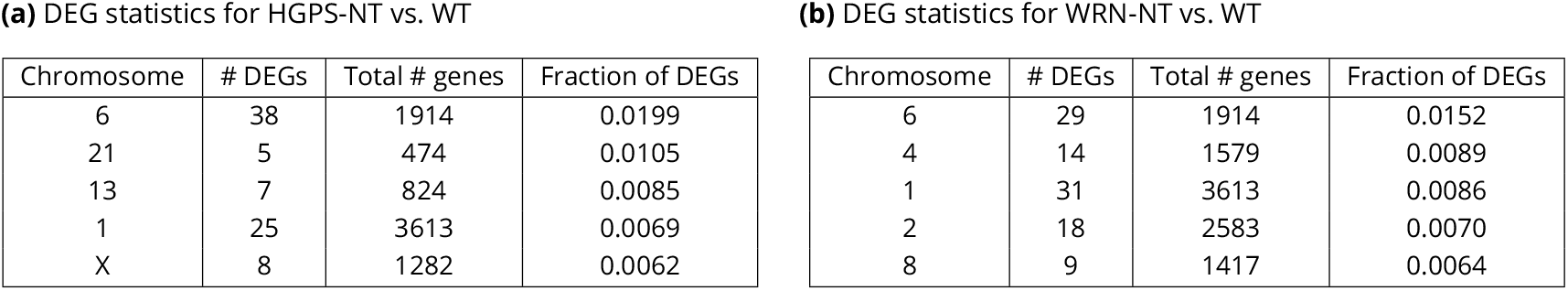
Differential gene expression analysis results **(a)** HGPS-NT vs. WT, and **(b)** WRN-NT vs. WT in L1 dataset. Number of differentially expressed genes (DEGs) in the top 200 hits, total number of genes, and the fraction of DEGs per chromosome, are listed. Entries are sorted by the fraction of DEGs. Top 5 results are shown here. Chromosome 6 has the largest number of fraction of DEGs in both cases.

**Figure 5.**
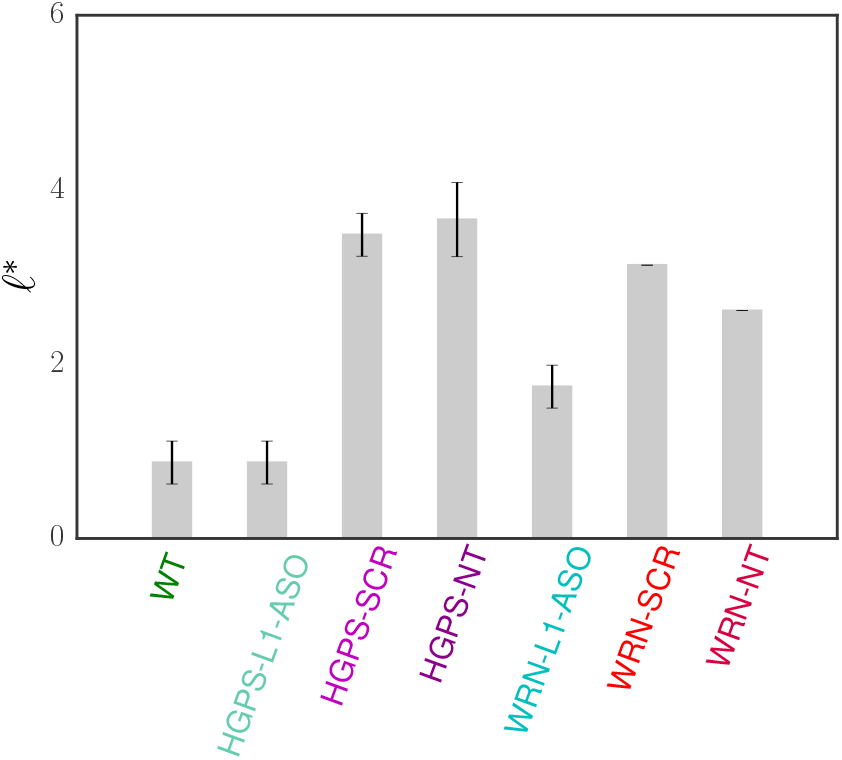
*Intra-chromosomal gene correlation length* (*ℓ*^*\**^) measured in LINE-1 dataset for chromosome 6. Error bars are computed from within group replicates. The figure illustrates that diseased samples exhibit a higher values of *ℓ*^*\**^ than the ASO treated/WT samples in chromosome 6.

These two observations confirm that *ℓ*^*\**^ not only captures state differences that are also evident in RNA-seq data but can also serve as a measure of changes in chromatin structure, which in turn influences gene expression. The increased correlation seen in both diseased and older samples likely reflects the degradation of heterochromatin, a process associated with aging and age-related disorders. This degradation leads to the unraveling of chromatin structure, exposing previously repressed DNA regions to transcription factors and activating nearby genes. Once activated, these genes can influence distal genes, resulting in longer correlation lengths across chromosomes. Importantly, these epigenetic changes do not affect all chromosomes equally.

### Clustering on intra-chromosomal gene correlation length yield optimal results

Given the hypothesis that the proposed measure captures the properties of chromatin, it remains to be shown whether the proposed methodology can help assess *how far* a treatment shifts a cell line from a diseased phenotype towards a healthy control group based on its chromatin state. To illustrate this aspect, ***Figure 6a*** presents the clustering of the experiments in the LINE-1 dataset (see Methods section Hierarchical clustering). The figure shows the clustering results from our proposed metric and compares it with those obtained from PCA and SVD using the same dataset (see Methods section Dimension reduction algorithms, ***Supplementary Figure S4a, b*** and c). As mentioned earlier, the analysis in the main paper is limited to genes within the HAT class. Additional results, including those for HAT+LAT and LAT-only classes, are provided in ***Supplementary Figure S3***. It is important to note that since *ℓ*^*\**^ utilizes replicate information to select genes for the HAT class, the emphasis should not be on the initial clustering of replicates. Instead, the focus should be on the second-order clustering, which captures the relationships between different cell states. The key objective here is to examine the order of merging across different cell states, highlighted in pink.

**Figure 6.**
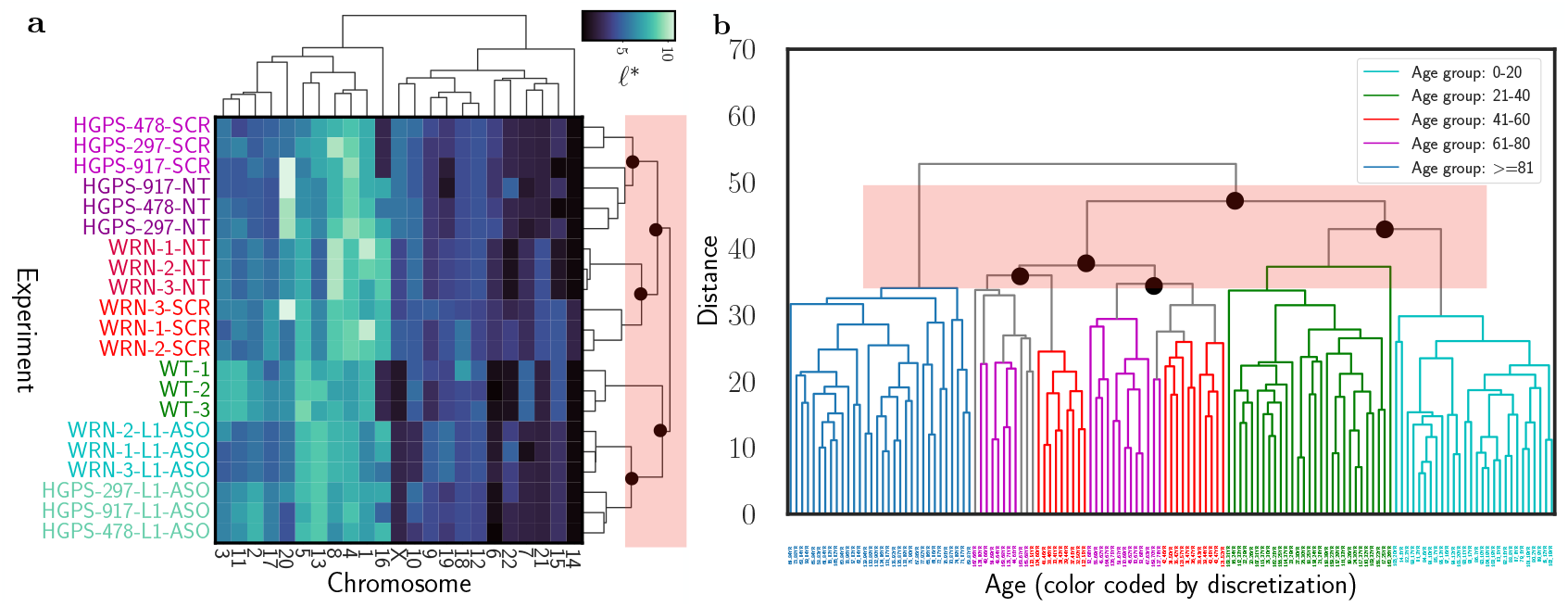
Hierarchical clustering of L1 dataset and Fleischer dataset. **(a)** Heat-map with dendrogram using the proposed *intra-chromosomal gene correlation length* (*ℓ*^*\**^) shows a clear clustering of the samples based on treatment and disease on the second level (highlighted by pink shade). **(b)** Hierarchical clustering of Fleischer dataset. Samples were bucketized into different age group and *ℓ*^*\**^ can cluster age groups in an expected order on the second level (highlighted by pink shade).

Recall that our proposed algorithm yields one value (*ℓ*^*\**^) per experiment for each chromosome. Each chromosome is represented along the abscissa of the heat-map in ***Figure 6a***. The figure demonstrates that the ASO treatment on LINE-1 moves the HGPS and Werner cell lines closer to the wild-type, indicating success as reported in ***Della Valle et al. (2022***). In contrast, the scrambled ASO treatment is less effective, clustering initially with its corresponding NT (no treatment) cell lines. Furthermore, the similarity between HGPS and WRN is highlighted through merging of their corresponding cell-lines (and variants) in both ASO and non-ASO settings.

A similar hierarchical clustering for the Fleischer aging dataset is shown in ***Figure 6b***. In this dataset, samples were grouped in the age ranges of 0 − 20, 21 − 40, 41 − 60, 61 − 80 and 81 − 100 to demonstrate trends in the correlation lengths over ages. Once again we observe that using *ℓ*^*\**^ leads to a robust clustering of the data. We observe that the younger age groups (0 − 20 and 21 − 40 age bracket) merge first. Concurrently, we see the middle age group merging together (the 41 − 60 age group with that of 61−80). It is noteworthy that within these age groups, there are two sub-clusters, which may be due to the variation in chronological ages versus cellular ages. The senior age group (≥ 81) merges with the rest of the cluster last, highlighting the significant differences in *ℓ*^*\**^ in older individuals. On the other hand, clustering performance using PCA (***Supplementary Figure S4d***) is considerably poorer, further reinforcing that our proposed metric outperforms these traditional linear analyses.

### Gene rank varies across samples and cell states

The *intra-chromosomal gene correlation length* (*ℓ*^*\**^) helps characterize the chromatin states by measuring whether State *A* is closer to State *B* or State *C* in a way that is sensitive to the influence of chromatin structure on gene expressions. Given the known effects of aging on chromatin, *ℓ*^*\**^ can also provide insights into the relative order of age of these different states - if *ℓ*^*\**^ for State *A* is less than that of State *B*, we can infer that State *A* is younger than State *B*. However, it does not measure how easily one state can transition into another.

To assess this, we need to measure the transition energy barrier, which quantifies how easily the system can transition from State *A* to State *B* compared to State *C*. This requires examining the data in a “relative” space and comparing the chromatin structure of each state. When epigenetic modifications alter the euchromatin domains (due to aging, disease, or treatment), the expression — and consequently the rank — of certain genes will increase, while others will decrease. In other words, as chromatin transits across different states, the corresponding chromatin structure influences the expressions of different genes, leading it to transit across different “rank” values across states. Previous analyses of transcriptomic data have used gene rank to study distribution patterns and Zipf’s law (***Ogasawara et al., 2003***; ***Furusawa and Kaneko, 2003***; ***Lazzardi et al., 2021***). However, in this work, we are interested in using changes in rank between states as a proxy of the relative changes between cell states through which we can build a transition matrix, modeling shifts from State *A* to State *B*.

***Figure 7a,b*** shows the curves for gene expression for chromosome 1 (a similar characteristic curve has been observed for other chromosomes), sorted by rank for each experiment in L1 and FL dataset respectively. The rank is obtained by sorting the genes by their expression in descending order. Thus, the gene that is the most highly expressed is rank-1. A curve collapse, similar to previous studies, across different experiments can be seen for each chromosome. The inset of ***Figure 7a*** shows an example for two genes, *IRF6* and *AK4*, which are both on chromosome 1. The blue circles show an increase in the rank position (i.e., a decrease in expression level) for *IRF6* following LINE-1 ASO treatment, suggesting downregulation by the diseased states. In contrast, the red circles show a decrease in rank position (i.e., an increase in expression level) for *AK4*, indicating upregulation by the diseased state. We demonstrate a similar trend in the Fleischer dataset. Thus, one can see that ‘nearby’ states have similar ranks (WT vs. ASO, age bracket 21-40 vs. 0-20), whereas a markedly different rank levels for significantly different states (WT vs. NT, age bracket 81+ vs. 0-20). The objective here, through this example, is to demonstrate that a transition across markedly different states would result in different rank positions and these changes are not in isolation, i.e., a rank change of gene *x* would affect the rank change of gene *y*. The choice of *IRF6* and *AK4* is not to show that there is a coregulation relationship between them; rather the opposite, that irrespective of the relationship between pairs of genes, there may be global reordering of gene ranks even if only one gene changes its rank. The magnitude of this global reordering depends on how different are the two cell states. Extending the idea presented in the inset figures that show the transition of ranks across states for two genes, we create a transition matrix comprising of all rank transitions made by all the genes. In the following section, we demonstrate how this transition matrix is then used to estimate the energy landscape associated with chromatin modifications, using the Arrhenius equations and thermodynamic principles.

**Figure 7.**
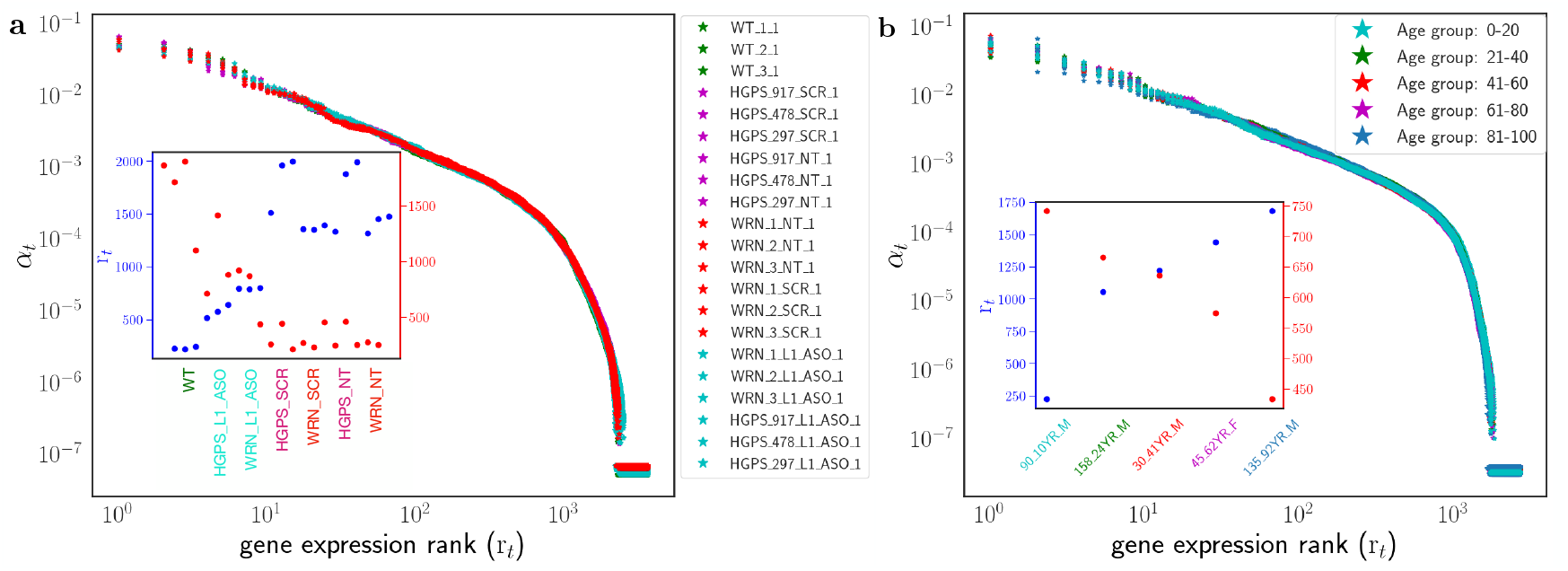
Gene expression (*α*_*t*_) as a function of rank (*r*_*t*_) for chromosome 1 in **(a)** LINE-1 dataset and **(b)** Fleischer dataset. The main plot shows that the normalized expression is conserved on a log-log scale while rank changes as a function of disease or treatment. The inset shows two example of how rank changes by experiment for two particular genes, *IRF6* (blue circles), and *AK4* (red circles). Pearson’s correlation coefficients and p-values are annotated on the figure. The samples along the x-axis in the inset are based on the rank of gene *IRF6* sorted from low to high.

### Aging and related disorders increases variability in chromatin energy landscape

In this section, we introduce the concept of a “chromatin energy landscape.” This landscape can be seen as an indirect measure of the reaction rate in a regulatory chemical process that drives the transition between two states. Using the Arrhenius equation, we estimate the energy barrier associated with changes in rank during this transition. Focusing on specific genes, such as those in the HAT class, and their rank transition data between two cell states, we calculate the energy landscape corresponding to the chemical process involved in transitioning from cell State *A* to cell State *B*. The detailed construction of this landscape is outlined in Methods section Chromatin energy landscape.

***Figure 8*** shows four energy landscapes for rank transitions between cellular states on chromosome 1. Additional examples can be found in ***Supplementary Figure S7*** and ***Supplementary Figure S8***. The abscissa in each landscape represents a distinct cellular state being compared (the comparand state), while the ordinate represents the same reference state across the landscapes. These landscapes provide two-dimensional visualizations of energy surfaces, where the intensity of each point corresponds to lower energy levels (disregarding color). The upper triangle of the heatmap represents the energy change for genes increasing in rank, whereas the lower triangle reflects the energy change for those decreasing in rank. The dispersion of points within each heatmap reflects the variance of states between the considered sample pairs. ***Figure 8a*** illustrates a nearly symmetric energy landscape for observed WT-WT transitions. For transitions within the same state (as shown in ***Figure 8a***), the energy change would ideally be zero, assuming identical replicates. However, slight variability among replicates results in a small spread along the diagonal. The lack of points distant from the diagonal suggests a high energy barrier, indicating a low transition rate for those rank pairs. For transitions involving jump across different states (***Figure 8b-d***), a larger dispersion along the diagonal is observed. Notably, ***Figure 8***b shows lesser dispersion than ***Figure 8c*** and ***Figure 8d***. This is in agreement with our observation when we apply *ℓ*^*\**^ on this dataset and the authors of ***Della Valle et al. (2022***) that LINE-1-treated ASO helps restore heterochromatin and brings cells closer to WT. Together, these findings suggest substantial changes in transcriptomic variance between states (representing aging), and the transition energy barrier offers a systematic means of quantifying these changes.

**Figure 8.**
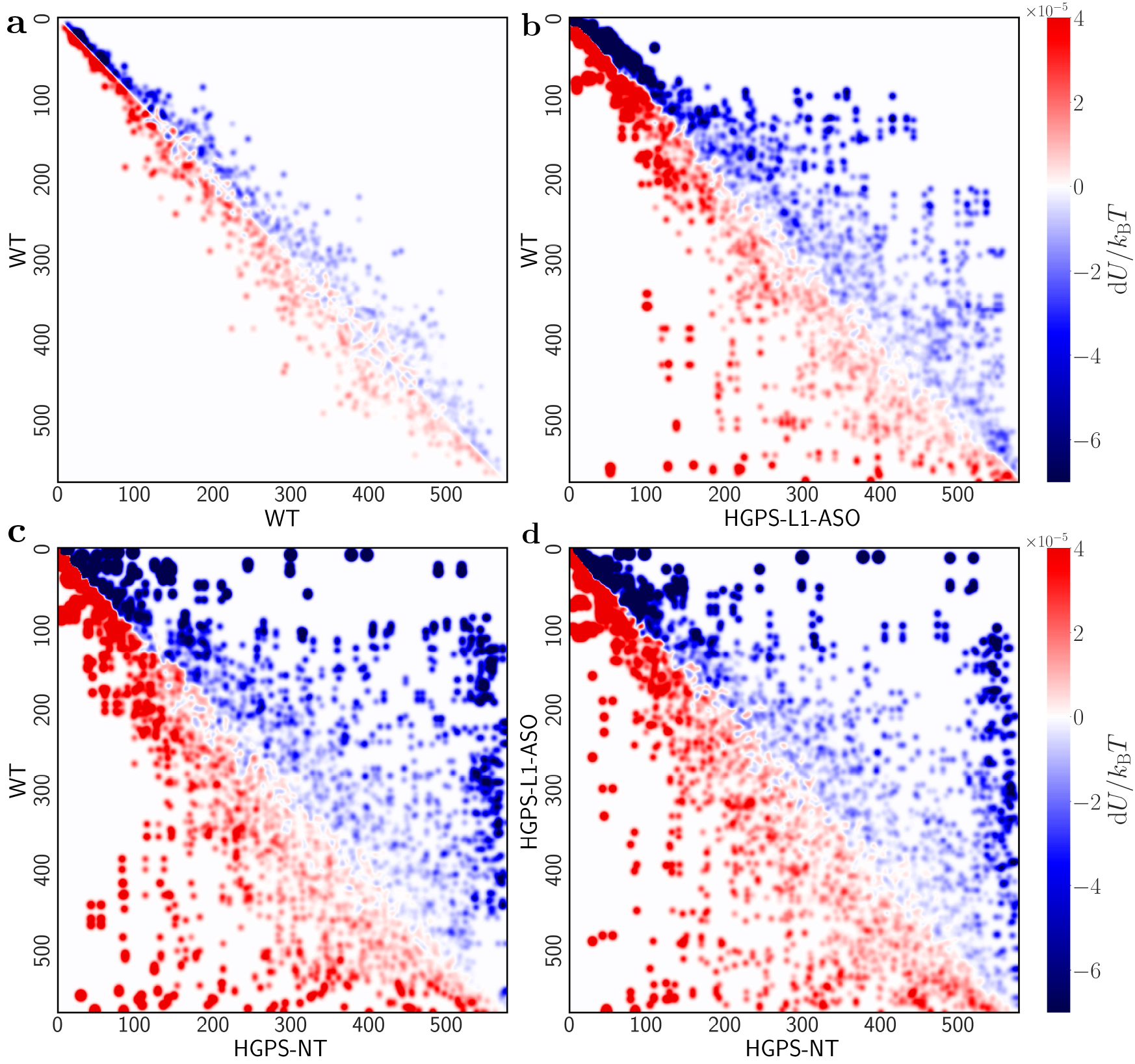
Chromatin energy landscape for chromosome 1 based on rank transitions and gene expression, applying the Arrhenius equation to the L1 data. The magnitude (intensity of each point) is based on the change in abundance for each rank change and replicate pairs given the two sample types in each sub-figure. Ordering the experimental classes by class(x) ⇒ class(y), the sub-figures correspond to: **(a)** WT ⇒ WT **(b)** HGPS-ASO ⇒ WT **(c)** HGPS-NT ⇒ WT and **(d)** HGPS-NT ⇒ HGPS-ASO. The upper triangle shows energy changes associated with genes moving up in rank, the lower triangle shows the energy change for movement down in rank. With the exception of panel **(a)**, the landscapes are not symmetric.

### High transition energy barrier associated with diseased/old cell states

We now introduce a quantitative measure that aggregates all the information from the 2D energy landscape, allowing us to quantify the transition energy barrier beyond visual inspection. To accomplish this, we begin with the 2D landscape shown in ***Figure 8*** and compute the total change in energy (Δ*U*) over all possible transitions between ranks. This is evaluated using the following double integral,

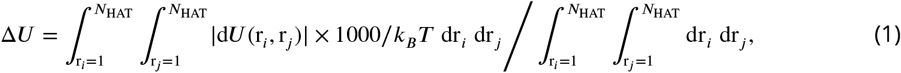

where the integral is performed over the rank space shown in ***Figure 8***.

***Figure 9a, b*** and c shows the corresponding transition energy barrier summed over all chromosomes for HGPS, WRN and Fleischer dataset respectively. The energy changes are plotted separately for genes that move up in rank (upper matrix in ***Figure 8***) and for those that move down in rank (lower matrix). The black curve (diff) in ***Figure 9*** shows the difference in Δ*U* between the upper and lower Δ curves. Observe that the transitions within the same states, such as WT ⇒ WT and Grp_1_20 ⇒ Grp_1_20, have a lower energy barrier (≈ 0) associated with them suggesting that it is energetically easier to transition within the same state. ***Figure 9a, b*** show that the energy required to transition from ASO-treated state to WT is lower compared to transitions between WT and diseased states (HGPS-NT ⇒ WT or WRN-NT ⇒ WT). This suggests that treatment with Line-1 ASO reduces the transition barrier for reverting from a diseased to a healthy state. While this is true for both the diseased states, the transition barrier is higher for WRN samples compared to HGPS. We perform the same analysis for the Fleischer data as well (***Figure 9c***), considering the younger age group (1 − 20) as the reference state. We also observe that a significant additional energy requirement to move from an old age group (≥ 80) to young, compared to the middle age-group and vice-versa.

**Figure 9.**
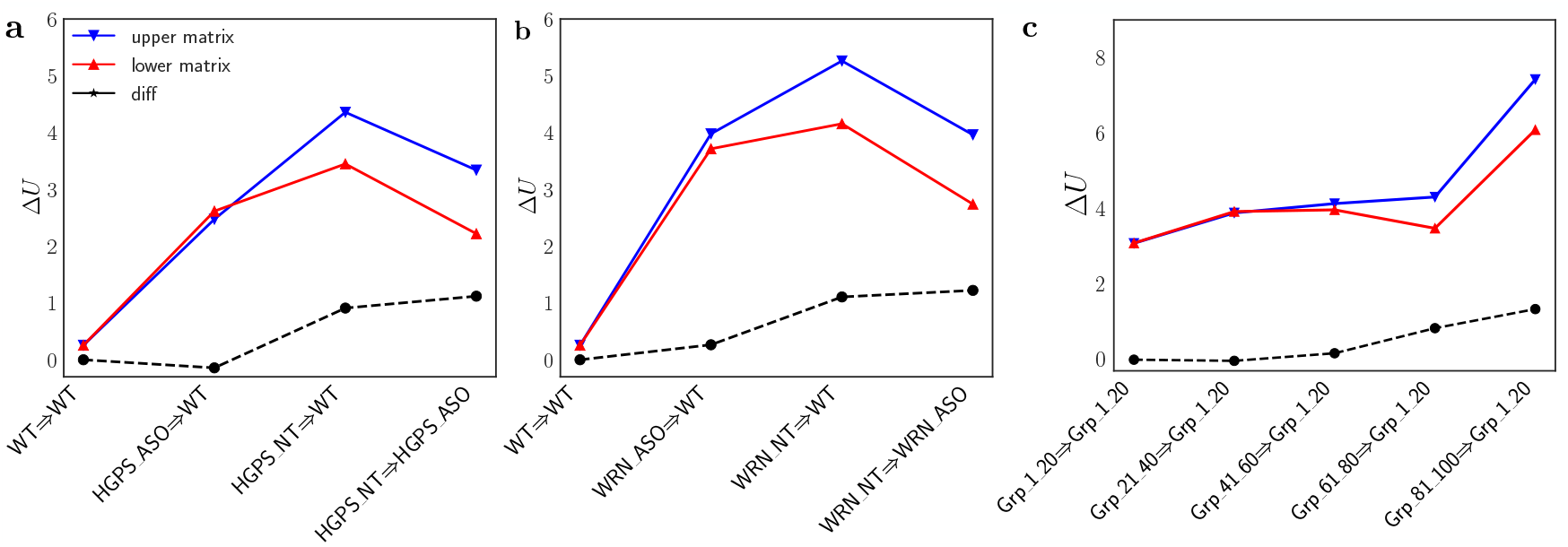
Chromatin transition energy barrier summarized across all chromosomes for transitions between samples up and down the ranks respectively using the integral expression (***Equation 1***). The markers with blue triangles shows the energy required to move up the ranks is relatively higher for diseased to WT/ASO transitions whereas WT to ASO transitions are equally probable in **(a)** L1 (HGPS) and **(b)** L1 (WRN). Moving from young to old state in **(c)** Fleischer dataset requires the most energy with higher variance compared to transitions from other age groups.

Interestingly, the two measures indicate the same value for WT ⇒ WT, HGPS-ASO ⇒ WT, with black curve ≈ 0. A non-zero value would indicate an *irreversible* nature of the additional cost to switch between the two states. In contrast, the irreversible energy component increases with age as well as in the untreated diseased state. These observations suggests a potential utility of this metric as an indicator of the changes in chromatin structure. The near zero difference for WT ⇒ WT and HGPS-ASO ⇒ WT transitions implies that these states have similar chromatin state, while the increasing irreversible component with age or in diseased states reflect more significant alterations in chromatin structure that would require more energy to reverse.

## Discussion

In this paper, we propose two macroscopic measures - intrachromosomal gene correlation length (*ℓ*^*\**^) and transition energy barrier. Both measures provide indirect insights into chromatin structure, offering supplementary perspectives that, together, deliver a more holistic view of cell states, particularly in the context of aging. To demonstrate efficacy of this work, we apply both these metric to aging and related syndromes datasets.

In the case of intrachromosomal gene correlation length, our results show a systematic increase in aggregated *ℓ*^*\**^ values with age. This increasing correlation length, *ℓ*^*\**^, appears to be consistent with the dysregulation associated with heterochromatin erosion, leading to ‘leaky’ genes and the introduction of low frequency signals. But are all chromosomes equally affected and should we expect an increase in *ℓ*^*\**^ for all chromosomes? To understand this, it is important to recall that chromatin is spatially organized, and this organization evolves during biological processes such as aging, differentiation, and development. Changes in the nuclear lamina during aging (or accelerated aging conditions like progeria) directly affect chromatin structure and, consequently, gene expression. Additionally, distinctions between constitutive and facultative heterochromatin, and their distribution during biological processes, play a role, as does the fact that gene expression itself can remodel or constrain chromatin structure. For example, replicative senescence, often associated with aging, is characterized by the formation of senescence-associated heterochromatin foci (SAHF), a type of facultative heterochromatin linked to specific nuclear features (***Hänzelmann et al., 2015***). Since SAHFs are not uniformly distributed across the genome, it is reasonable to expect differences in *ℓ*^*\**^ across chromosomes (***Supplementary Figure S11***). While in the case of Fleischer dataset, we did observe a systematic increase of *ℓ*^*\**^ over age, the two syndrome studies (HGPS and WRN) did not demonstrate this pattern. Rather, they exhibited strong trends in chromosomes that also happened to be most differentially expressed.

In the Line-1 dataset, chromosomes 6, X and 16 are the top-3 chromosomes in terms of proportional increases in *ℓ*^*\**^ with respect to WT in HGPS-NT, while in WRN-NT, chromosomes 16, 6 and X show the largest changes (*ℓ*^*\**^ for LINE-1 and Fleischer dataset is available in the Additional Files category). This suggests that heterochromatin erosion may be most pronounced on these three chromosomes. Previous studies have shown that the largest cluster of histone genes (*∼* 80%), HIST1, is located on chromosome 6 and *FOXO3* ‘interactome’ on chromosome 6 is a chromatin domain that defines an aging hub (***Donlon et al., 2017***). A rare abnormality on chromosome 16 has been associated with disease progression and increased nuclear localization of WRN (***Slupianek et al., 2011***; ***Yanagiya et al., 2020***). Differential gene expression analysis also indicates that chromosome 6 has the highest proportion of differentially expressed genes for NT vs. WT states for both syndromes (***Table 2***). In the case of Fleischer as well, if we compare each age group with the 0-20 age bracket as the reference, we still observe that chromosome 6 is the top chromosome in terms of proportional increase. However, in the case of the Fleischer dataset, we do not observe any strong statistically significant continuous trend of individual chromosomes over all age brackets. Rather, as shown in ***Figure 4***, we see a global trend that arises due to the increased statistical power with aggregation of all chromosomes. This further bolsters the belief that aging appears to be a systematic loss of regulation genome-wide and occurs incrementally. This further highlights the difference between pathological and natural aging in the case of fibroblast cell lines.

In this paper, we additionally show how *ℓ*^*\**^ could be used as a feature in a downstream application, e.g., clustering cell states. We show that because of its ability to capture non-linear spatial effects (due to the use of autocorrelation on bucketized distance correlation) arising from structural changes to chromatin, it surpasses the performance of traditional linear methods such as PCA and SVD. Furthermore, the proposed *intra-chromosomal gene correlation length* models and captures the underlying aging mechanism and consequently is richer in terms of semantics it represents. It is also interesting to note that in ***Figure 6a*** chromosome 8 merge with 4 and then 1. Chromosomes 1 and 8 harbor genes associated with HGPS and WRN syndrome, respectively. Previous studies have revealed that these two syndromes share a common pathological mechanism associated with progerin expression, an abnormal variant of lamin A protein (***Kang et al., 2021***). The protein coding gene *LMNA* is located on chromosome 1. Furthermore, studies such as ***Murano (1995***); ***Puca et al. (2001***); ***Liao et al. (2021***); ***Yeh et al. (2022***) have shown that chromosome 4 has genes that are associated with longevity and genes related to musculoskeletal disorders.

In the case of the transition energy barrier, our findings reveal that the barrier energy increases as the differences in the cell states increase. This trend aligns with existing published literature and further supports the notion that homeostatic regulating mechanisms deteriorate with aging. Consequently, this method is reflective of the concept of “buffer capacity”, which is indicative of cell resilience. As the cell states under examination represent various stages of aging, the trend of the transition energy barrier offers a transcriptomic view of declining cellular resilience across these states.

An additional insight into chromatin structure changes emerges from the concept of “irreversibility” observed in the energy landscape. Recall that using the change in relative abundance of individual genes between pairs of cell states, we derived a transition energy barrier between the two states shown as the chromatin energy landscape (***Figure 8***). This landscape illustrates the energy changes associated with genes upregulation and downregulated between the two states as a result of alterations in chromatin state. If the total energy integrated over the upregulated genes (represented in the lower triangular matrix of the landscape) is precisely equal to the total energy integrated over the downregulated genes (upper triangular matrix), the landscape is symmetric, indicating that the process is thermodynamically “reversible”. For instance, consider a chromosome with a set of genes in euchromatin and another set of genes in heterochromatin. Simply repositioning all nucleosomes from the heterochromatic regions to the euchromatic regions has an energy cost. For a perfectly symmetric landscape, the net energy change cost for up/downregulation of genes resulting from nucleosomes reshuffling would be identical. However, if chromatin rearrangement due to epigenetic modifications leads to net addition or removal of nucleosomes, the two energies would not be equal, with the difference representing an irreversible energy cost. Specifically, if a disease state reduces nucleosome numbers on DNA, adding ‘missing’ nucleosomes incurs additional energy costs, which will apply to genes that move up in rank (down in expression level).

This irreversible energy component is demonstrated by the dashed black line in ***Figure 9***. Notably, for transitions within the same group, such as WT⇒ WT and Grp_1_20 ⇒ Grp_1_20, the energy cost for upregulation and downregulation is nearly identical (difference ≈ 0). In contrast, the irreversible energy component increases with increasing age leading to a non-zero difference. The same is true when comparing no treatment to WT or ASO. These observations are consistent with the view that Progeria, Werner syndrome (L1), and aging (FL) all lead to a loss of nucleosomes, and that treating or correcting these conditions requires energy to replace the missing nucleosomes.

Furthermore ***Figure 9a,b*** suggests that heterochromatin erosion in HGPS is more severe than WRN for the cell lines studied. The total energy cost is similar for untreated WRN and HGPS (HGPS-NT ⇒ WT and WRN-NT ⇒ WT blue points in ***Figure 9a,b***). ASO treatment leads to a greater reduction in total energy cost for HGPS than WRN (HGPS-ASO ⇒ WT and WRN-ASO ⇒ WT blue points in ***Figure 9a,b***. These findings underscore the need for better intervention and treatment designs for Werner syndrome, as ASO treatment proves highly efficient for HGPS.

Taken together, our results are fully consistent with the idea that chromatin structure constrains gene expression. Traditional studies of cellular reprogramming and disease treatments rely on a relatively small number of biomarkers to indicate whether the treatment or reprogramming succeeded. These biomarkers are microscopic probes. The macroscopic approaches proposed in this work provide quantitative measures of how much a cell state changes in response to disease or treatment. Our approach can be further strengthened by adding multimodal data, including ATAC-seq/ChIP-seq/Hi-C data, to identify microscopic changes which may lead to the discovery of new biomarkers.

## Methods

### Data pre-processing

In this study, we utilize two publicly accessible human whole genome bulk RNA-seq datasets. These datasets are useful for testing our hypothesis regarding the large-scale disruption of gene expression caused by alterations in chromatin structure, as well as for estimating the transition energy barrier related to the epigenetic differences between cell states.

1. LINE-1 (L1): The Long Interspersed Nuclear Element-1 (LINE-1) dataset (***Della Valle et al., 2022***) contains human fibroblast samples for 7 cell lines, each containing 3 replicates. ***Table 1*** provide additional details about this dataset. The GEO accession ID is GSE198675.
2. Fleischer (FL): The Fleischer dataset (***Fleischer et al., 2018***) contains human fibroblast cells from 143 donors of different ages ranging from newborns to 96 years. Amongst the 143 donors, there are 10 donors with HGPS. We remove these samples from our dataset to avoid misleading analysis and conclusion. The GEO accession ID is GSE113957.

In the case of LINE-1, we additionally mapped the ensembl ids to human genome GRCh38.p13 using the BioMart - Ensembl tool to obtain the chromosome information. Genes that do not express themselves in any experiment/cell line (i.e., their expression values are 0s in the entire dataset) are dropped. Each experiment/cell line in the dataset, containing the fragments per kilo-base of transcript per million mapped reads (FPKM) values is then normalized by the sum to obtain the fraction of each gene contribution to total expression in a cell. This is denoted as *α*_*t*_ in the main text.

### Intra-chromosomal gene correlation length

The computation of the intra-chromosomal gene correlation length is outlined in Algorithm 1. Given the division of the genes into the HAT and LAT classes, we quantify the positional dependence of gene expression by calculating the autocorrelation function of normalized RNA-seq data (*α*_*t*_). We begin with binning the pairwise expression values based on the chromosomal positions of the genes with each bin containing transcript abundance (**x, y**) for pairs of genes, and calculate the distance correlation for each bin. The distance correlation 𝒞 (Δ*ℓ*) coefficient is given by the

#### Algorithm 1

Intra-Chromosomal Gene Correlation Length Calculation Per Chromosome

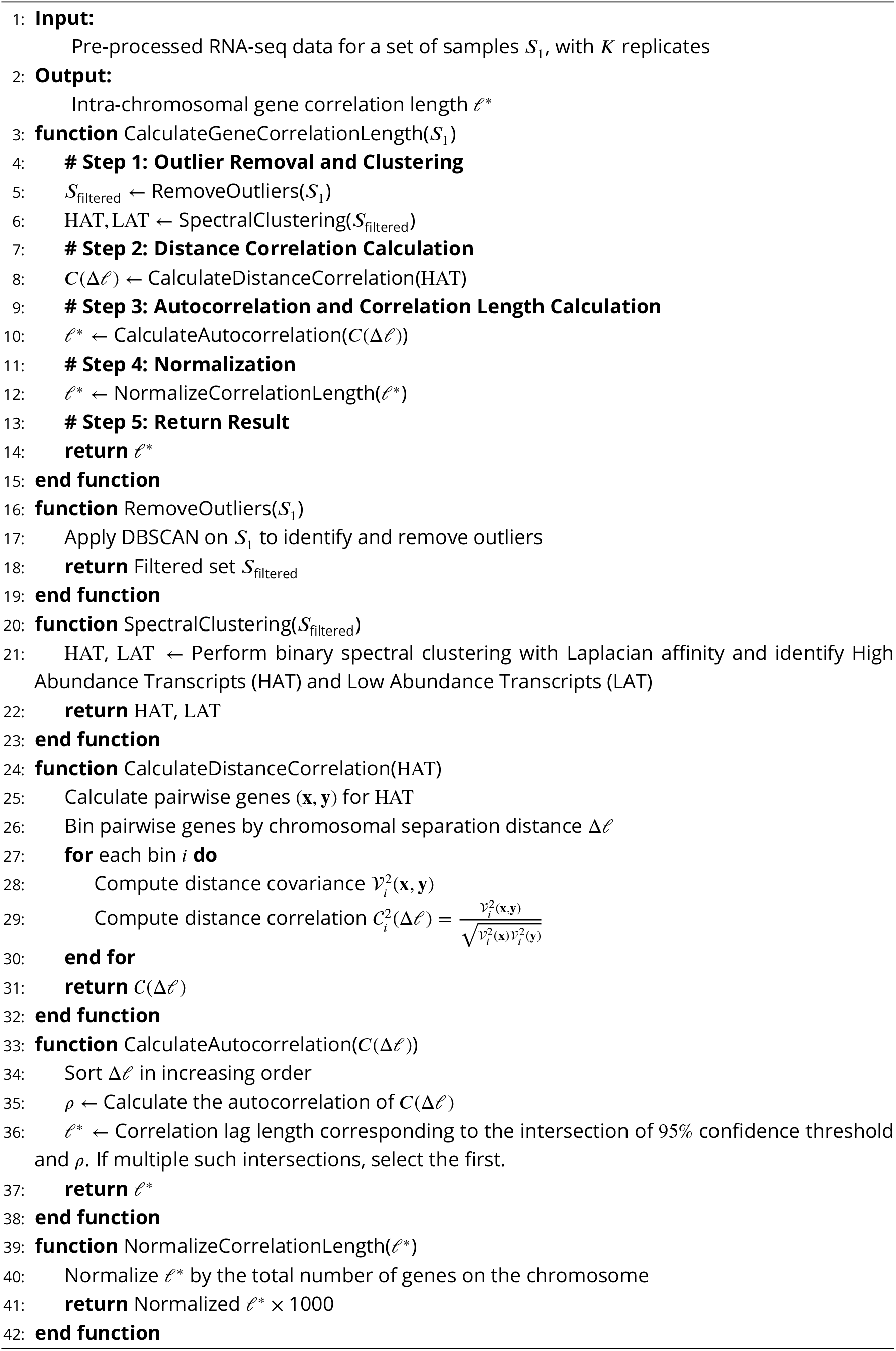

following mathematical formulation

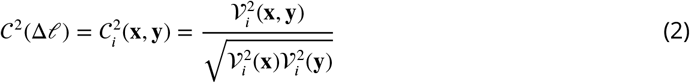

where each pair in bin *i* is separated by Δ*ℓ* number of genes between them and finally, empirical distance covariance denoted by 𝒱 can be written in terms of centered Euclidean distances 𝒟,

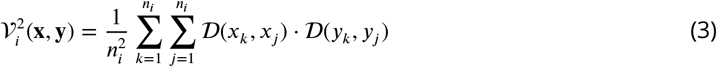

We further perform the autocorrelation of 𝒞 (Δ*ℓ*) which measures the relationship of a variable with lagged values of itself and hence, provide with a measure to obtain the length scale. Performing autocorrelation is useful to capture the scale of distance correlation decay till the point of sharp increase for high (Δ*ℓ*). We then measure the correlation length for each chromosome as the point where the gene-gene autocorrelation falls below the 95% confidence limit, and the correlation length at this intersection point is then normalized with the number of genes on the corresponding chromosome. This normalized value is then defined as the *intra-chromosomal gene correlation length ℓ*^*\**^. 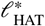 calculated from the HAT class used as *ℓ*^*\**^ unless specified otherwise explicitly. Note that fine-tuning the gamma parameter for spectral clustering is essential, especially for downstream analysis. The gamma parameter is reflective of the input data complexity. If the input data are relatively easier (e.g., LINE-1) to separate, say using the PCA, lower gamma (around 1) is preferred. For more complex data like Fleischer, we set gamma to higher values (around 3). Additional discussion on this subject can be found in the Supplementary.

### Dimension reduction algorithms

We use PCA and SVD as exemplars of dimension reduction algorithms. The input to these algorithms is the normalized RNA-seq data - same as the one to the distance correlation module explained in the previous section. We use ‘scikit-learn’ packages to implement PCA and SVD.

### Hierarchical clustering

The derived features (*ℓ*^*\**^, or PCA, SVD) are fed to hierarchical clustering to merge replicates and cell-lines. The linkage algorithm used in ‘Average’ and the metric to calculate distance is ‘city block’ (a.k.a Manhattan distance). In fact, it is to be noted that all the distances measured in this paper are using city block distance. Advantage of using city block distance over Euclidean distance is shown in ***Supplementary Figure S9***. We use ‘clustermap’ function from *seaborn* Python package which in turn uses SciPy.

### Manhattan distance on gene expression and intra-chromosomal gene correlation length

To calculate the similarity distance shown in ***Figure 3***, we use the feature matrix for *ℓ*^*\**^ and normalized abundance transcript (*α*_*t*_) to construct the cluster distance matrix 𝒞 _p_(*ℓ*^*\**^) (red curve) and 𝒞 _p_(*α*_*t*_) (blue curve) respectively. The distance is calculated using the Manhattan distance between each pair of samples using wildtype as a reference to compare the similarity between two experimental phenotypes. The mean across different chromosomes can be used to plot the average cluster distance along with standard deviation, calculated across replicates, as an error bar.

### Statistical testing

The Mann-Kendall Trend Test is implemented using original_test() in pyMannKendall package (***Hussain and Mahmud, 2019***) and the outputs are reported. The Mann-Whitney U rank test is implemented using mannwhitneyu() in SciPy package and the outputs are reported.

### Differential gene expression analysis

Raw counts of RNA-seq from the LINE-1 dataset were used. Genes with low counts across one or more experimental conditions were excluded. Subsequently, the PyDESeq2 package (***Muzellec et al., 2022***), a Python-based tool for bulk RNA-seq differential expression analysis, was employed to conduct the differential analysis. We compared the differentially expressed genes (DEGs) between the three experimental groups (HGPS-NT vs. WT and WRN-NT vs. WT) with a p-adjusted cutoff of 0.0001, and found that the diseased samples exhibited 3,153 DEGs. Among these, 1,535 were upregulated, while 1,618 were downregulated.

### Chromatin energy landscape

The computation of the energy landscape is outlined in Algorithm 2.The input to the algorithm consists of RNA-seq data from two samples, *S*_1_ and *S*_2_, each representing different cell states that we are interested in comparing. Each sample *S*_*j*_ (*j* ∈ {1, 2}) is composed of *K* biological replicates, denoted 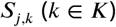, where 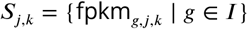 represents the FPKM values for all transcripts *g* in the set *I*.

The algorithm begins by sorting the transcripts for each replicate within *S*_1_ and *S*_2_ in descending order based on their FPKM values. Each transcript is then assigned a rank 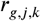, where *g* represents the transcript, *j* denotes the sample (either *S*_1_ or *S*_2_), and *k* indicates the replicate. The transcript with the highest abundance is assigned rank 1, while the one with the lowest abundance is given the lowest rank, corresponding to the total number of transcripts.

After ranking, the algorithm identifies the transitions of transcripts between ranks across all replicate pairs from *S*_1_ to *S*_2_. For each transcript *g*, the algorithm tracks how the rank 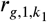 in a replicate *k*_1_ of *S*_1_ transitions to rank 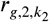 in a replicate *k*_2_ of *S*_2_. These transitions are accumulated into a transition matrix 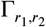, which counts how many times a transcript transitions from rank *r*_1_ in *S*_1_ to rank *r*_2_ in *S*_2_.

The transition matrix is then normalized to produce a probability matrix *p*(*r*_1_, *r*_2_), calculated using the expression:

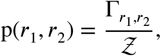

where 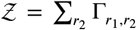 is the sum of all transitions for a given *r*_1_, ensuring that the probabilities are properly normalized. The resulting probability matrix reflects the likelihood of a transcript transitioning between any two ranks across the different experimental conditions.

To derive the activation energy landscape, the probability matrix is transformed using the Arrhenius equation, where the change in activation energy d*U*(*r*_1_, *r*_2_) is given by:

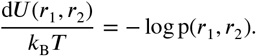

In this equation, *k*_B_*T* is the thermal energy factor, and the logarithmic transformation converts the probability matrix into an energy landscape, where lower energy indicates a higher probability of transition between ranks. Before applying the logarithm, the matrix is adjusted to ensure non-negative values by offsetting it such that the minimum value is zero. After the transformation, the matrix is re-centered. A Gaussian filter is then applied to smooth the data and interpolate across any sparse regions in the transition matrix, enhancing the clarity of the resulting energy landscape. The final energy landscape is visualized as a heatmap, as shown in ***Figure 9***, with additional sensitivity analysis on the Gaussian filter parameters provided in ***Supplementary Figure S10***.

## Acknowledgements

We want to acknowledge helpful discussions with Benjamin Yang, Erik Navarro, Rebekah Brooks, and Simone Bianco.

### Algorithm 2

Calculation of Activation Energy Landscape Per Chromosome

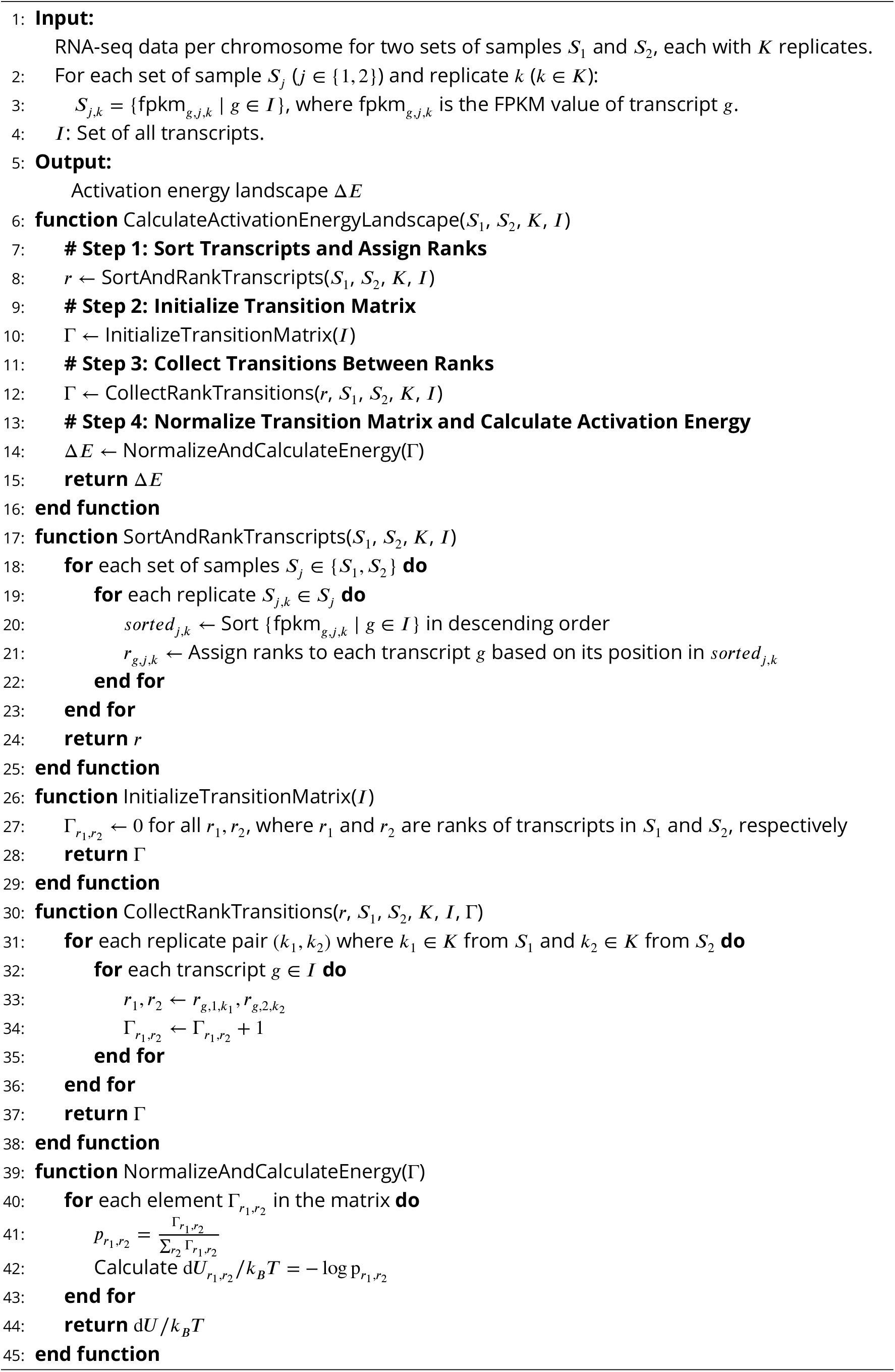

## Code and data availability

Python scripts and data for this manuscript are available on GitHub: https://github.com/altoslabs/papers-2025-rnaseq-chrom-aging

## Supplementary Material

**Figure S1.**
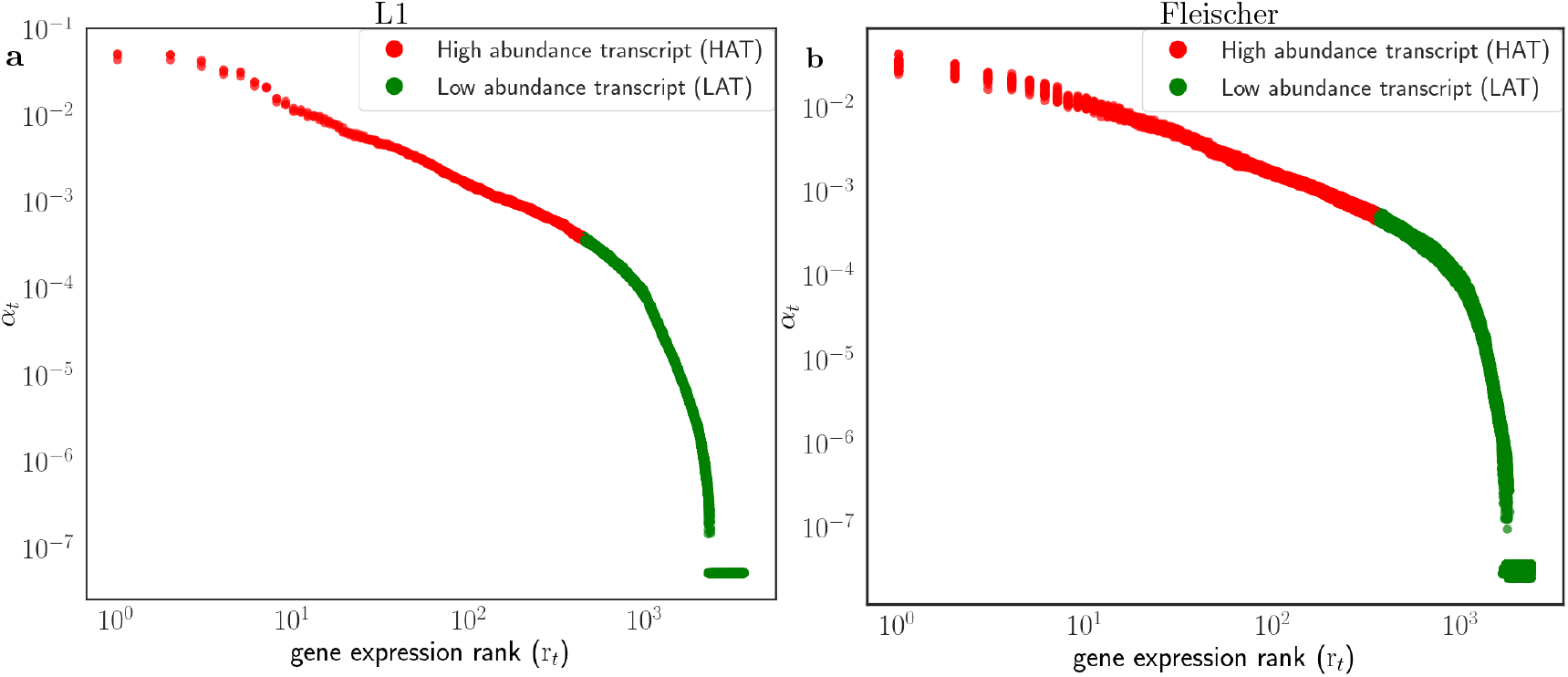
An unbiased spectral clustering procedure is used to segregate all gene based on their expression values (*α*_*t*_) into one of two groups: HAT and LAT represented by red and green markers respectively. The plots depict clustering results for **(a)** WT with 3 replicates in L1 dataset, and **(b)** samples from the 0 − 20 age group, both corresponding to chromosome 1. The x-axis represents the rank of genes, determined by sorting their expression values in descending order.

**Figure S2.**
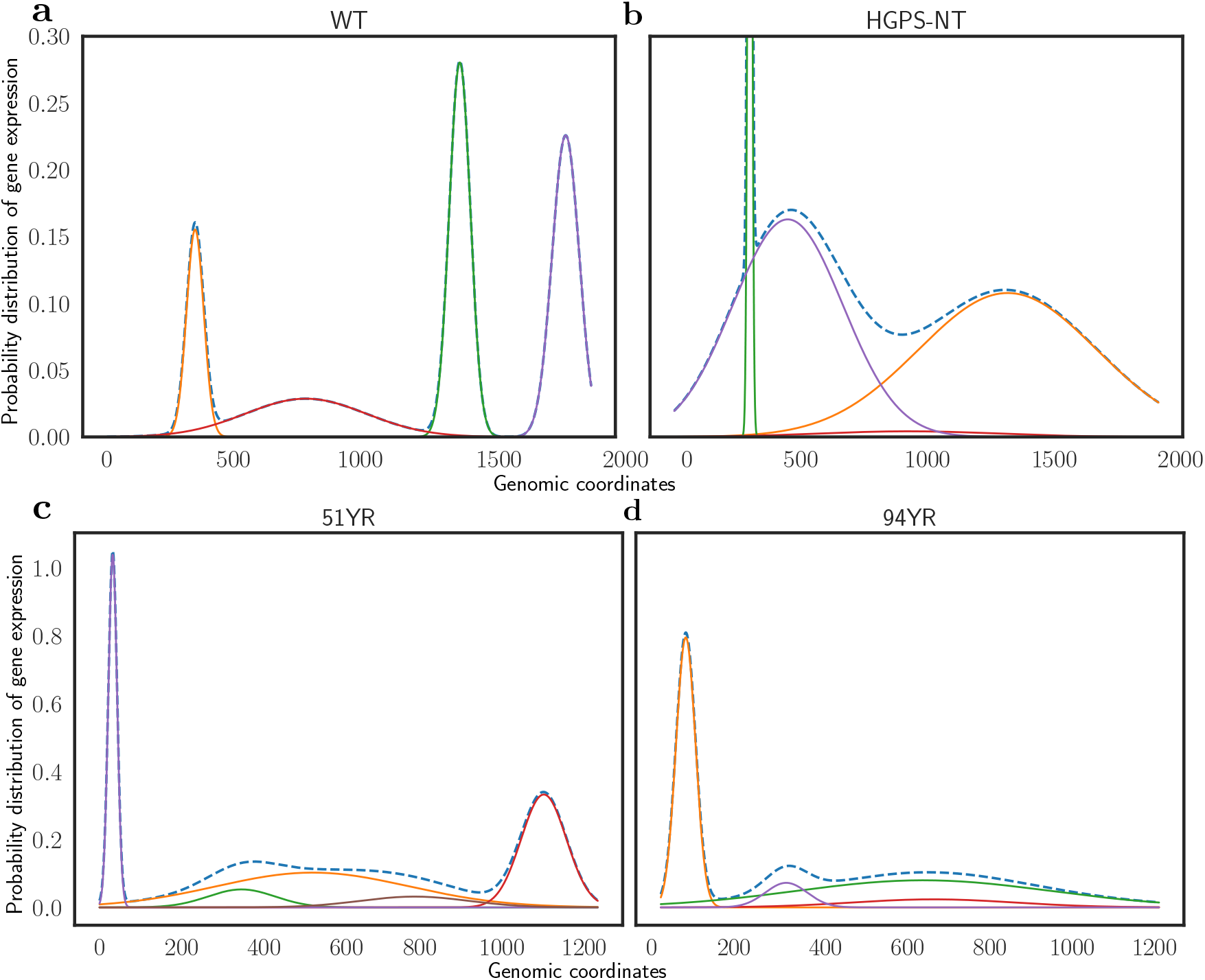
Comparison of gene expression activity for Chromosome 6 is shown between healthy (WT) samples in sub-figure (a) and diseased (HGPS) samples in sub-figure (b) from the LINE-1 dataset, as well as between a 51-year-old sample (sub-figure c) and a 94-year-old sample (sub-figure d) from the Fleischer dataset. The x-axis in each subplot represent the genomic coordinates of the chromosome. One can see sharply defined distribution characteristics for the healthier and “younger” samples and more flatter distributions spanning multiple coordinates for the diseased and “older” samples. The distributions are obtained by fitting Gaussian Mixture Model and using Dirichlet process to estimate the number of components. The flatter distribution spanning multiple coordinates indicate increased activation of “nearby” genes w.r.t target intended genes during the RNA-polymerase transcription. The sharper distribution observed in the healthier/younger population leads to a conclusion of a relatively intact regulatory mechanism that is able to more precisely activate the target intended genes.

**Figure S3.**
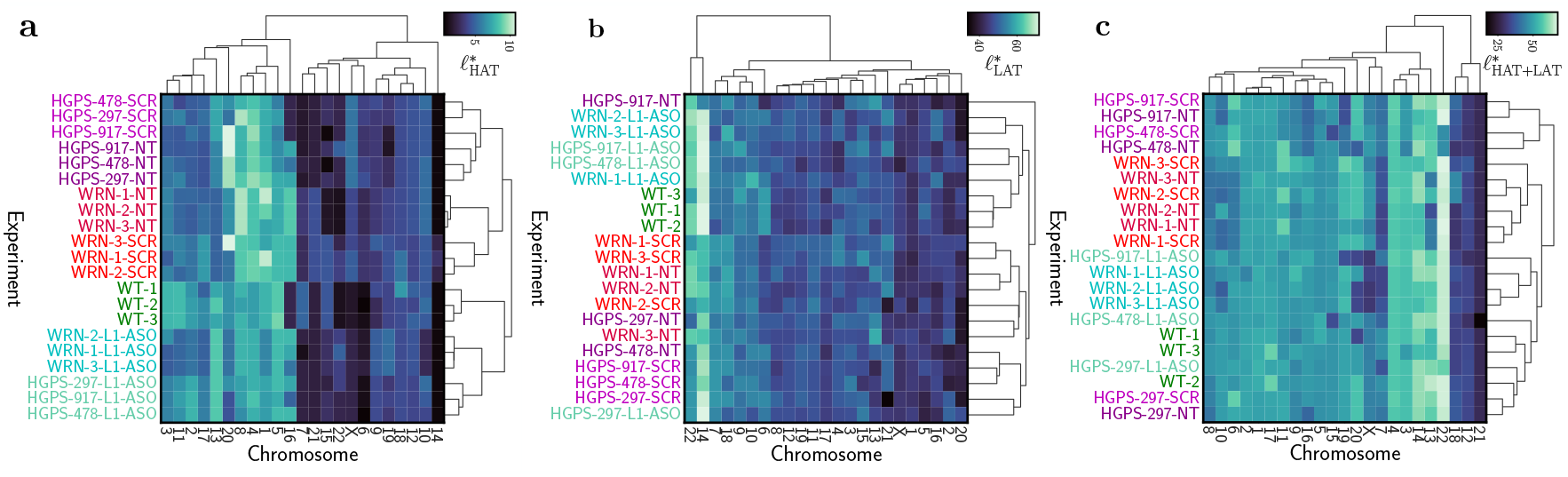
Comparison of hierarchical clustering using *intra-chromosomal gene correlation length* (*ℓ*^*\**^) in different scenarios for L1 dataset: **(a)** considering only HAT genes in the **(b)** considering only LAT genes **(c)** considering all genes. Clustering using HAT genes shown in **(a)** yields optimal results.

**Figure S4.**
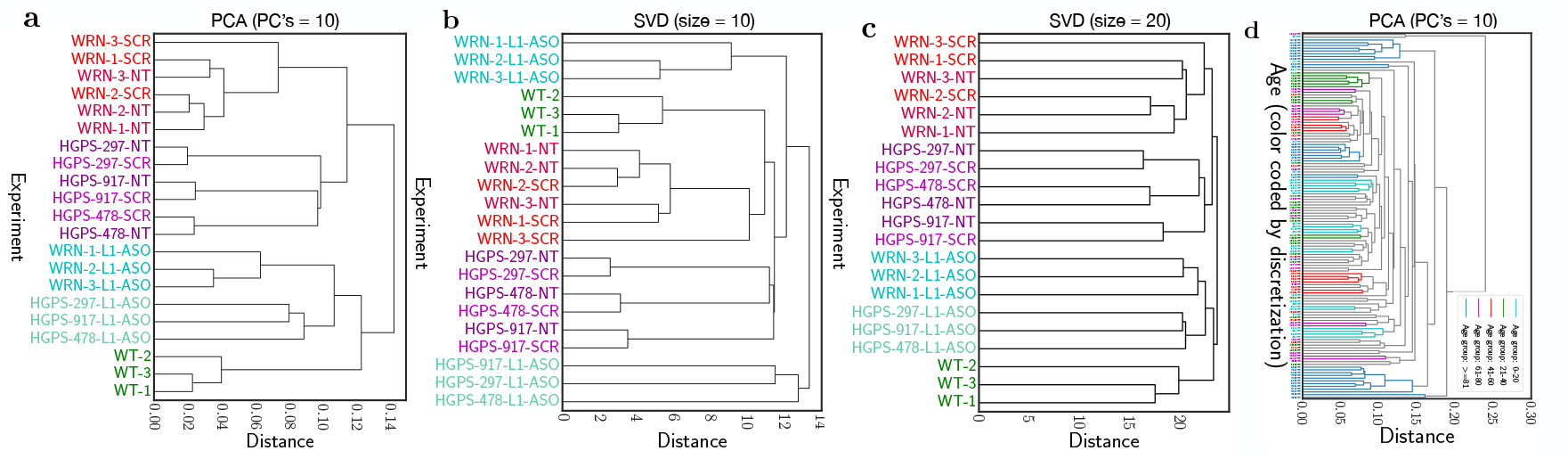
Hierarchical clustering results using SVD with **(a)** size = 10 and **(b)** size = 20 in FL dataset. Clustering using SVD yields poor results. The x-tick labels are color coded with cyan for 0 − 20 age group, green for 21 − 40, red for 41 − 60, magenta for 61 − 80 and blue for ≥ 81. The grey colored dendrogram lines represent instances where samples from different age groups are merged.

**Figure S5.**
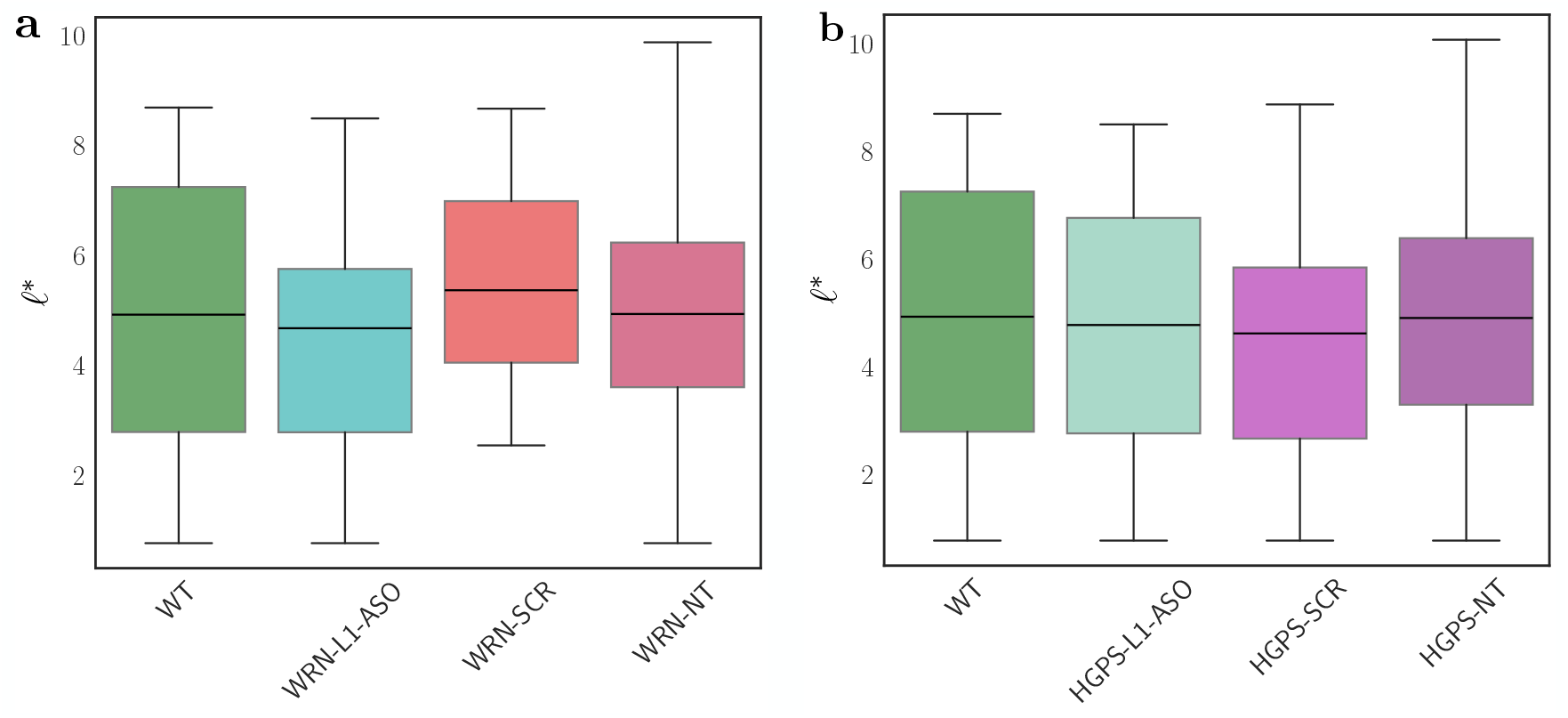
Box plots showing variation of *intra-chromosomal gene correlation length* (*ℓ*^*\**^) across different chromosomes. No significant differences were observed between untreated or scrambled-treated HGPS samples and those treated with Line-1 ASO, as well as WT samples, suggesting an alternative epigenetic modification mechanism in pathological aging. The line in each box plot represents the mean rather than the median.

**Figure S6.**
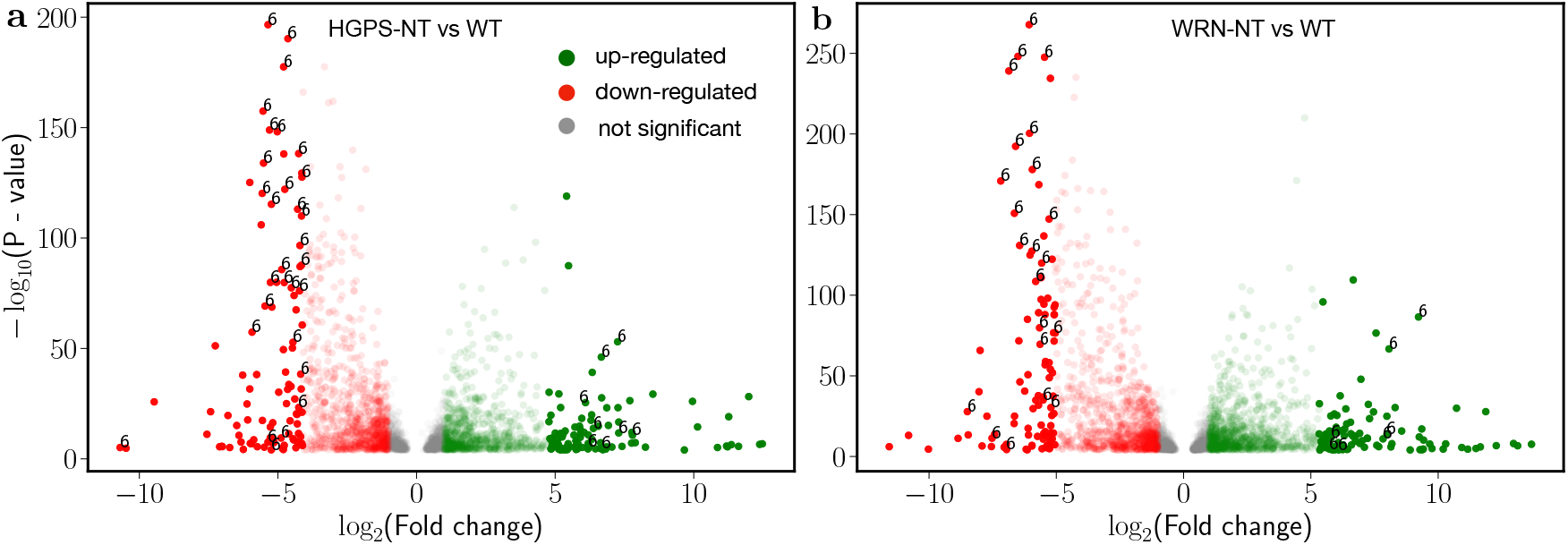
Differential gene expression analysis for three experimental conditions **(a)** HGPS-NT vs. WT and **(b)** WRN-NT vs. WT using PyDESeq2 to identify differentially expressed genes. The dark colored markers show the top 200 hits based on the log_2_ fold change out of which 38 (HGPS-NT vs. WT) and 29 (WRN-NT vs. WT) are found on chromosome 6 and annotated in the figure.

**Figure S7.**
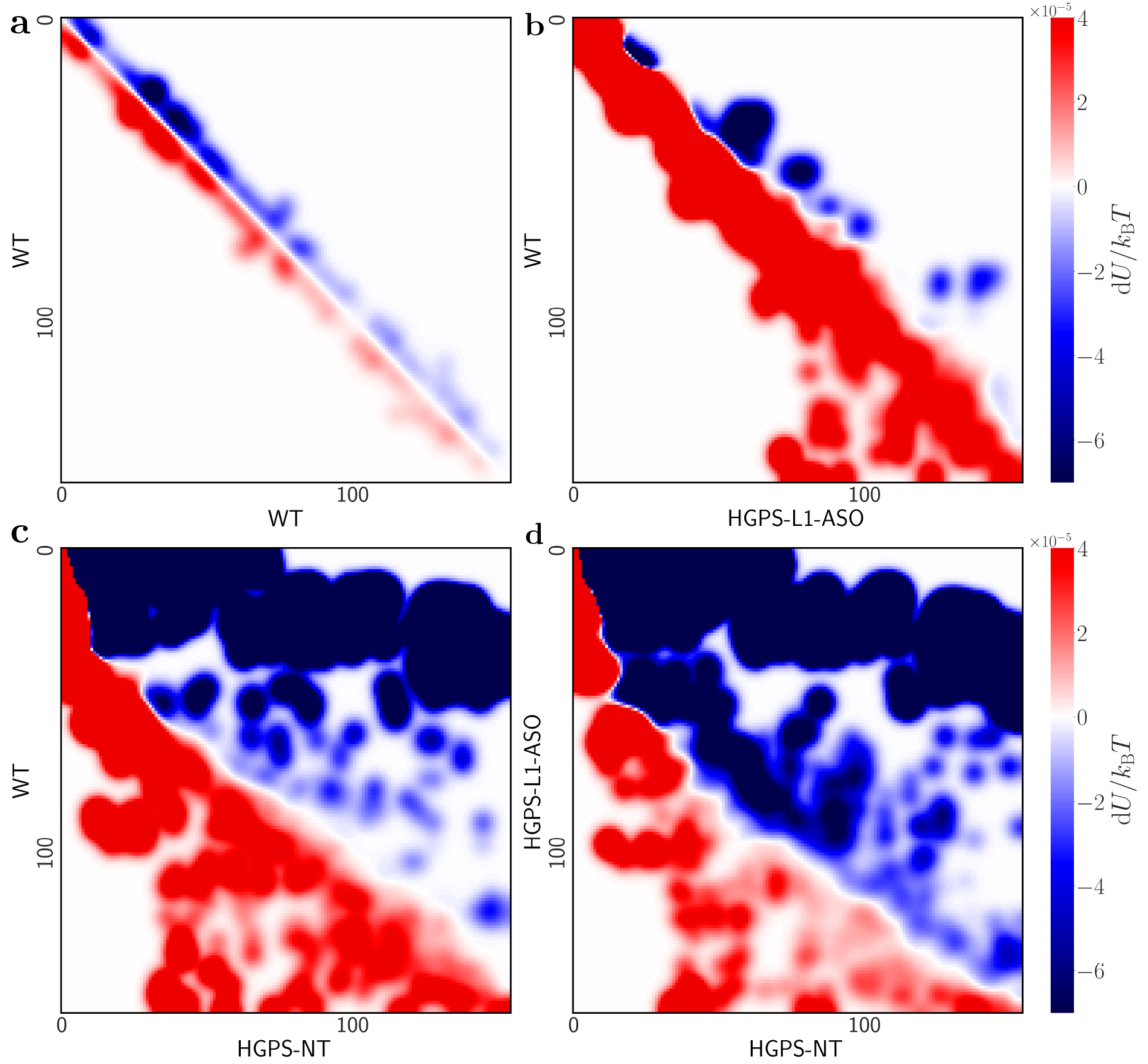
Chromatin energy landscape for chromosome 6 in the case of HGPS for L1 dataset, similar to Fig. 8 in main text. The energy landscape for all other chromosomes look similar.

**Figure S8.**
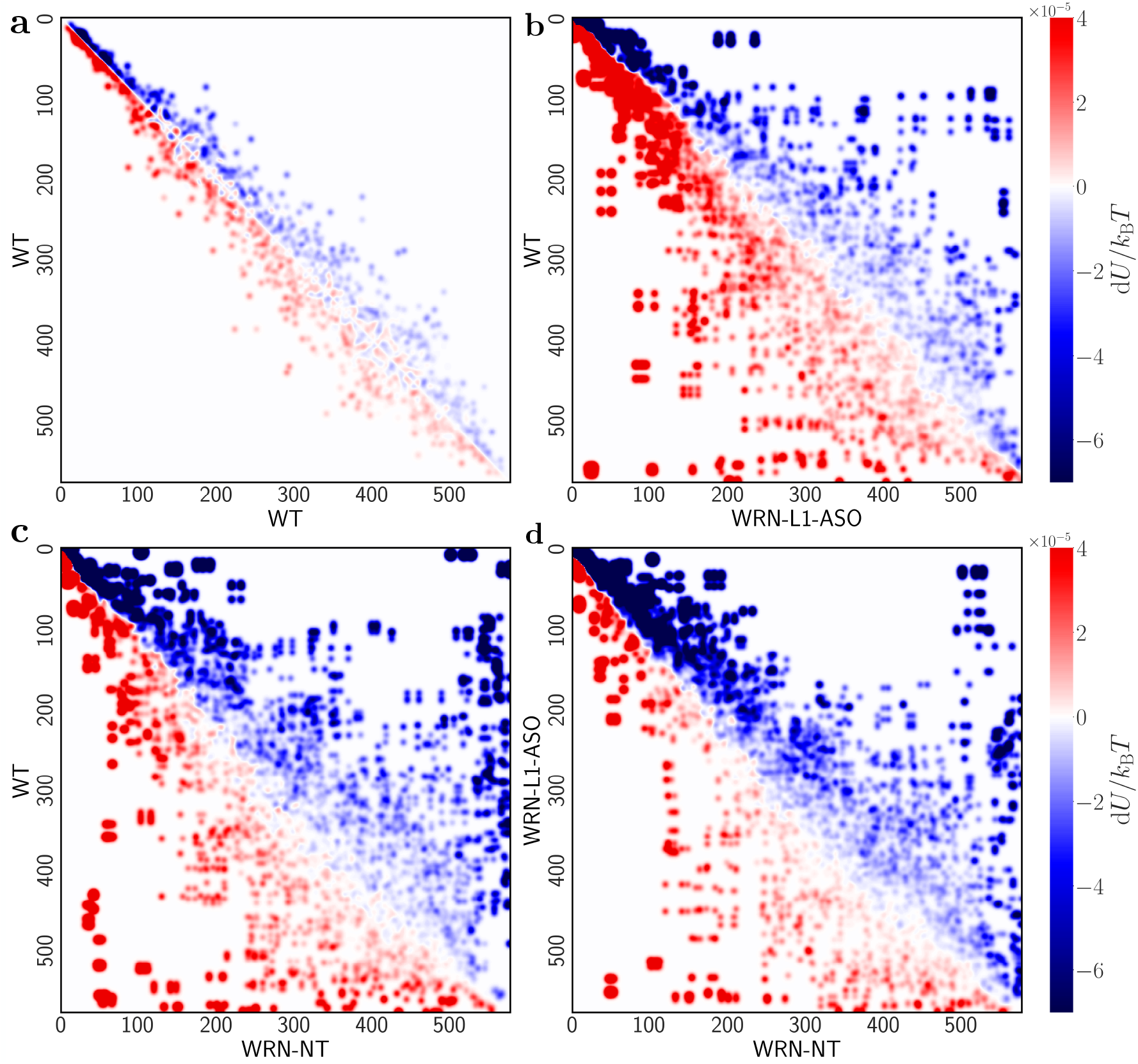
Chromatin energy landscape for chromosome 1 in the case of WRN for L1 dataset, similar to Fig. 8 in main text. The energy landscapes for all other chromosomes look similar.

**Figure S9.**
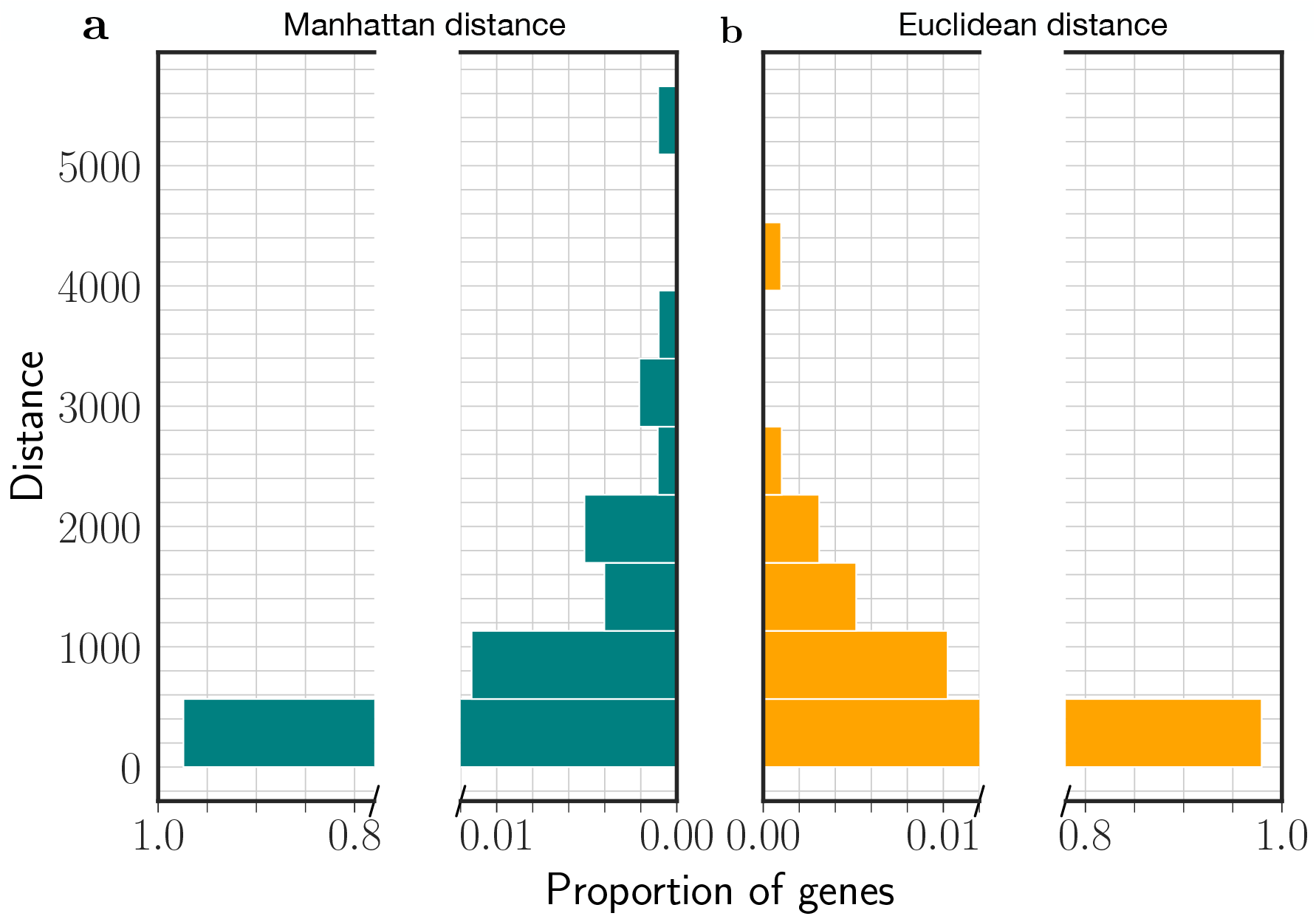
**(a)** Manhattan distance (city block) **(b)** Euclidean distance comparison when used in hierarchical clustering. One can observe the range of values that Manhattan distance uses in larger than the Euclidean distance. This leads to more effective discrimination between nearby points.

**Figure S10.**
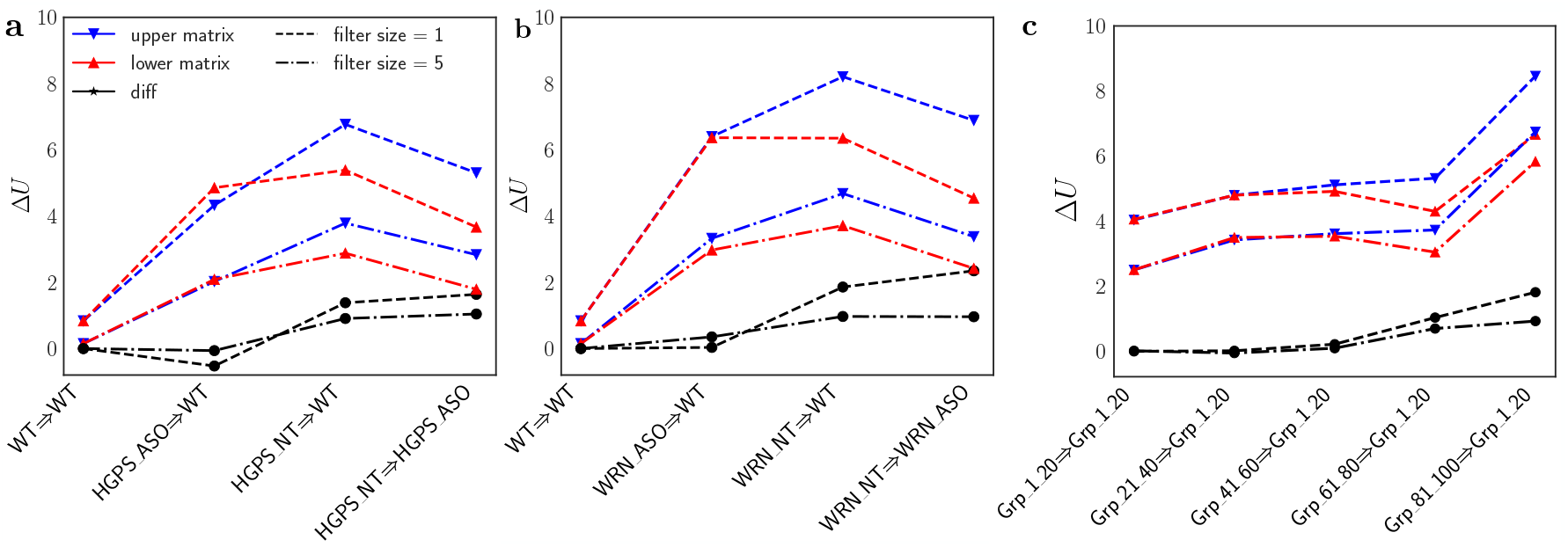
Chromatin transition energy barrier summarized across all chromosomes for transitions between samples up and down the ranks respectively for different sizes of Gaussian filter **(a)** L1 (HGPS), **(b)** L1 (WRN) and **(c)** Fleischer dataset. The trend is similar to Fig. 9 (filter size = 3) in the main text suggesting that the results are robust irrespective of the choice of filter size.

**Figure S11.**
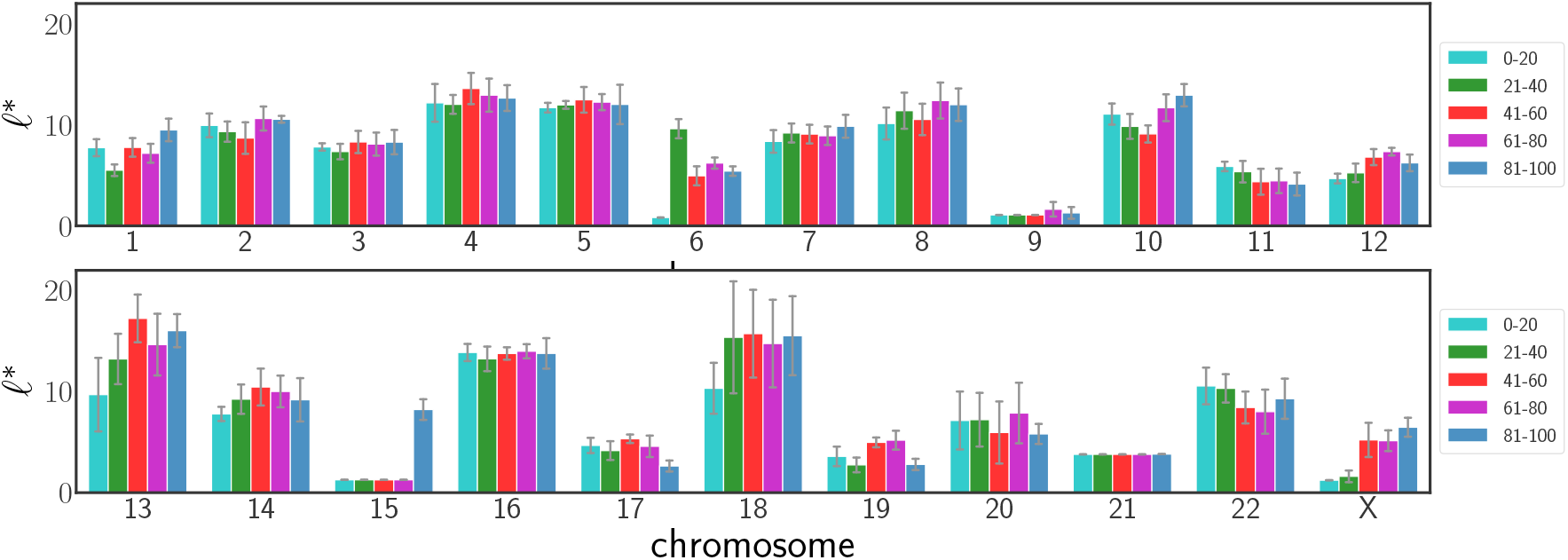
Chromosomal variation of *ℓ*^*\**^ for Fleischer dataset. Error bars represent the standard deviation across different samples within each age group.

## Effects of *γ* in spectral clustering

One of the important steps in the pipeline is categorizing genes into HAT and LAT classes. The algorithm we employ is Spectral Clustering with Manhattan distance affinity. An important aspect of the spectral clustering is the parameter *γ. γ* can be thought to be as one compensating for the complexity of the data. By complexity, we refer to how easy is it to distinguish different cell lines/states. This can be measured by calculating the Silhouette score on the PCA landscape. For a relatively well-separated landscape, we recommend *γ* = 1. For more complicated scenarios, we recommend trying a range between 2.5 and 3.5.

This brings us to an important question: How does *γ* affect the performance?

As we increase *γ*, the boundary that discriminates High Abundance Transcripts (HAT) from the Low Abundance Transcripts (LAT) shift towards higher expression values. In other words the precision of HAT class increases with increase in *γ* and correspondingly there is a decrease in the recall. Note, that our final *ℓ*^*\**^ value is calculated based only on transcripts in HAT. Similarly decrease in *γ*, moves the boundary towards the lower expression values, and increases the recall and decreases the precision of transcripts in the HAT class.

Increased precision and decreased recall result in leaky genes being discarded. Thus, the strength in the increasing trend of *ℓ*^*\**^ due to aging reduces as one increases *γ*. However, because of the high precision in the selected genes, there is an increase in the pipeline’s ability to distinguish different samples. Hence, with an increase in *γ*, we also observe an increase in the accuracy of clustering (reduced misclustering of samples in dendrogram).

Similarly, increased recall and reduced precision, adds too many noisy transcript to the system, corrupting the clustering ability. However, since we are including lot more genes and possibly covering almost all of the leaky transcripts, we get a more accurate trend of *ℓ*^*\**^. In extreme case, one can turn off the denoiser module and measure the trend but the resulting clustering using *ℓ*^*\**^ will be poor.

